# Sex-specific frontal cortical circuit mechanisms mediating fear extinction

**DOI:** 10.64898/2026.04.10.717763

**Authors:** Kourtney Graham, Justice Pomeroy-Tuck, Grace K. O’Brien, Lacey Kent-Webber, Anna L. Drillen, Luke Coddington, Xinyu Zhao, Erik B. Bloss

## Abstract

Strong evidence suggests synaptic plasticity is the critical cellular mechanism underlying learning and memory. Although the forms of synaptic plasticity used by different circuits vary, a widespread presumption is that the male and female brain has evolved to use the same form of plasticity within the same circuits during learning. We used complimentary approaches to determine how synaptic plasticity within the mouse frontal cortex supports extinction of associative memories. Here, we show that both male and female mice have similar ensemble dynamics in excitatory infralimbic cortical neurons during learning. However, activity in amygdala-projecting neurons was required for extinction memories only in male mice. Likewise, only male mice showed evidence for structural synaptic remodeling and clustering of dendritic spines on infralimbic-amygdala projection neurons. Projection-specific deletion of the glutamate receptor subunit GRIN2B blocked synaptic plasticity and impaired extinction memory in male but not female mice. These distinct mechanisms could be leveraged for precise therapies for mental health conditions relative to the present one-size-fits-all approach.

---

The frontal cortex has long been considered a top-down control center for coordinating behaviors^1^. Past work has shown a crucial role for frontal cortical circuits in rule learning and implementation, particularly when task contingencies change^2,3^. Mechanistic experiments in rodents have used associative fear learning and extinction paradigms to produce competing context-dependent task contingencies. In this paradigm, extinction learning and memory formation requires a network of interconnected cortical circuits including the amygdala, thalamus, and frontal cortex^4–6^. Neurons in the infralimbic cortex (IL) are thought to consolidate extinction learning, as IL activity is required during extinction learning for later extinction memory retrieval. Although past work has shown evidence of synaptic plasticity in frontal cortex during extinction learning^7^, the cellular logic, nature of the synaptic plasticity, and causal relationship to behavioral control has remained unclear.

Here, we find that neurons in the mouse infralimbic cortex show dynamic within-session and across-session representations of extinction-relevant cues during extinction learning and at extinction memory retrieval. Chemogenetic neural silencing during extinction learning revealed that activity in IL-to-basolateral amygdala (IL-to-BLA) projection neurons is required for subsequent extinction memory retrieval. Extinction learning coincided with structural synaptic plasticity and synaptic clustering on IL-to-BLA neurons, suggesting a rapid form of rewiring of cortical circuits during learning. Consistent with a causal role for synapse plasticity, deletion of the N-methyl-D-aspartate receptor (NMDAR) subunit GRIN2B from IL-to-BLA neurons coordinately impaired learning-related synaptic structural plasticity and performance during extinction learning and extinction memory retrieval. Surprisingly, the requirement for neuronal activity, remodeling of synaptic size and local dendritic clustering, and the dependence on GRIN2B in IL-to-BLA neurons was absent in female mice, suggesting distinct mechanisms mediate the same cognitive processes across male and female mice.

## IL excitatory neuron ensemble dynamics during extinction learning and memory retrieval

Mice rapidly form associative memories between conditioning tones and brief footshocks that present as increased immobility in response to the tone and can be extinguished in a context-dependent manner by sessions in which the tones are presented without a footshock in a distinct context (12 tones/extinction learning session, **Extended Data Figure 1A**). Persistent long-term extinction memory retrieval required two extinction sessions (**Extended Data Figure 1B**). Although male mice show greater rates of tone-dependent immobility compared to females during the first session, behavioral responses are indistinguishable between male and female mice during the second session, during extinction memory probe trials in the extinction context, and in the original associative context (**Extended Data Figure 1C**).

The infralimbic cortex (IL) has been implicated in the formation and retrieval of extinction memories^8–13^. To establish the patterns of IL excitatory neuron activity during extinction learning and during extinction memory retrieval, we expressed GCaMP6f in IL excitatory neurons and recorded neuronal activity in the form of intracellular Ca^2+^ transients (**Figure 1A-B**). Past work found increased IL neuron activity at tone onset as a corollary of extinction memory retrieval strength^11^, so we focused on the recruitment of tone-onset responsive activity within and across extinction learning sessions. In male and female mice (n=9 each), approximately 14% of IL excitatory neurons (n=179) were classified as tone-onset responsive (see **Methods**) during the first three tones in the first extinction learning session. Ca^2+^ transient amplitudes in these neurons decreased over subsequent tone presentations and were absent by the end of the session (**Extended Data Figure 2A-B**). This decreased activity occurred alongside the emergence of a new subpopulation of neurons that showed strong tone-onset responses at the end of extinction learning session 1 (i.e., during the last three tones, n= 116 cells, **Extended Data Figure 2A-B**). The same flip-flop between early tone-responsive and late tone-responsive populations was evident during the second extinction learning session and during extinction memory retrieval (**Figure 1D-E, Extended Data Figure 2C-D**). These dynamics are not present in the absence of learning during the pre-tone periods of either of extinction learning session **(Extended Data Figure 2E-G**). PCA trajectories show that extinction learning session 2 ensembles had the largest path length (**Extended Data Figure 3A**), suggesting that the evolution of ensemble dynamics is distinctly different during the second extinction learning session.

**Figure 1.**
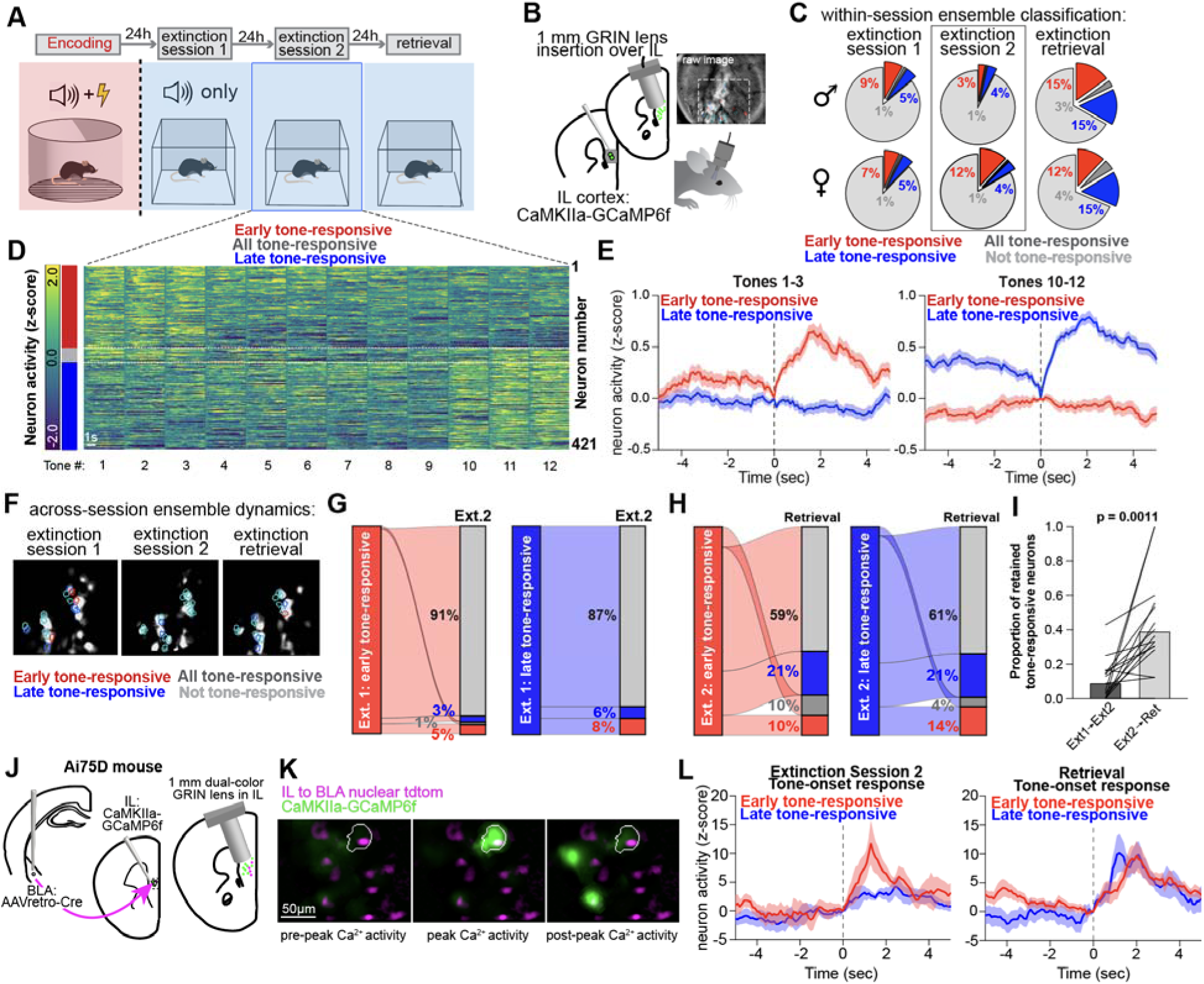
Dynamic IL excitatory neuron activity patterns within and across extinction sessions and at retrieval. **A**) Schematic of the behavioral paradigm used for context-dependent encoding, extinction and retrieval of extinction memories. **B**) CaMKIIa-GCaMP6f was injected into IL followed by a 1mm GRIN lens implant to image Ca^2+^ transients in individual excitatory neurons in IL during extinction and retrieval sessions. **C**) Male and female mice have similar proportions of tone-onset responsive neurons within extinction session 1 (n=1135 neurons from 9 male mice; n=1196 neurons from 9 female mice) and retrieval (n=928 neurons from 9 male mice; n=968 neurons from 9 female mice). Female mice have a greater proportion of tone responsive neurons than males during the early tone presentations on extinction session 2 (Fischer’s exact test, p<0.0001, n=1903 neurons from 9 male mice; n=1683 neurons from 9 female mice.) **D**) Heat map of individual neuron responses (plotted as normalized z-scores) for the first 5 seconds after tone onset within extinction session 2 (see **Extended Data** Figure 2 for extinction session 1 and retrieval). Neurons were classified as responsive to tone presentations early in the session (tones 1-3, n=255 neurons from18 mice), across all tones (n= 21 neurons from 18 mice), and to tone presentations late in the session (tones 10-12, n=145 neurons from 18 mice). **E**) Mean neural activity traces (shown as z-scores) from neurons in these classes during tones presented early in the extinction session (left) and from the same neurons during tones presented late in th extinction session (right). **F**) Images showing the spatially intermingled organization of neurons that were registered and classified across all three behavior sessions. **G**) Sankey plots showing the population dynamics of excitatory neurons across extinction sessions according to their tone-responsive activity show that neurons identified as tone-responsive during extinction session 1 were rarely classified as tone-responsive during extinction session 2. **H-I**) The proportion of tone-responsive cells identified from extinction session 2 that were also tone-responsive during extinction memory retrieval increased relative to that proportion across extinction sessions 1 and 2 (within-animal paired t-test, p=0.0011, n=18 mice). **J)** Dual color imaging enabled the identification of IL-to-BLA projection neurons in Ai75D mice, which express a Cre-dependent nuclear tdTomato reporter. **K)** Co-registration of the nuclear IL-to-BLA tdTomato signal with calcium transients during tone presentation show that IL-to-BLA neurons have tone-specific activity. **L)** Mean activity traces of IL-to-BLA neurons identified as early or late tone-responding reveal increased tone-responsiveness early on extinction day 2 (n=9 early tone-responsive neurons, n=8 late tone-responsive neurons, both taken from 5 mice) and at both early (n=20 neurons from 5 mice) and late (n=17 neurons from 5 mice) tones during extinction retrieval.

To establish the across-session dynamics of individual IL neurons, we tracked individual neurons that could be registered across extinction learning and extinction memory retrieval sessions (**Figure 1F**). The fraction of neurons that retained tone-onset responsivity across extinction learning sessions was only 10±3% (**Figure 1G**), which is consistent with the low fraction of neurons with persistent task-relevant representations across days in hippocampus^14^ and parietal cortex^15^. Male mice retained significantly fewer tone-responsive neurons from extinction learning session 1 to extinction learning session 2 than females (**Extended Data Figure 2E**, t-test, p=0.045), suggesting that males had a larger turnover of the IL ensemble of tone-responsive neurons between extinction learning sessions. Interestingly, there was a higher retention rate of tone responsive neurons between the second extinction learning session and extinction memory retrieval, with 37±7% of tone-responsive neurons maintained across memory retrieval when compared to 9±3% between the extinction learning sessions (**Figure 1H-I**, paired t-test, p=0.0011, n=18), suggesting stability in ensemble activity across memory consolidation. Collectively, these results show an emerging sex difference in IL neuron activity dynamics in which males have a higher turnover rate of tone-responsive cells between extinction learning sessions suggesting that males have more plasticity occurring during the learning phase, but both sexes maintain greater stability in ensemble dynamics during the memory consolidation phase.

IL projections to the BLA have previously been implicated in extinction learning and extinction memory retrieval^16,17^, but how these neurons participate in tone-responsive ensembles remains unknown. To establish these features, in a subset of mice (n=5 mice) we injected a retrograde AAV-Cre-recombinase virus into the BLA of Ai75D mice which express Cre-dependent nuclear tdTomato. In combination with AAV-CaMKII-GCaMP6f injections into IL, we used dual-color GRIN lens imaging to assign dual-labeled neurons as IL-to-BLA+ **(Figure 1J-K**). Registration of the tdTomato signal with the Ca^2+^ transient activity show that these IL-to-BLA projection neurons are recruited and highly active early during extinction learning session 2 (z-scored peak mean amplitudes of 11.8±3.6 and 4.1±2.6 for early and late tone-responsive neurons) and at extinction memory retrieval (z-scored peak mean amplitudes of 9.6±2.4 and 10.1± 3.2 for early and late tone-responsive neurons; **Figure 1L).** Together, these results show that IL-to-BLA neurons robustly encode tone signals during extinction learning and extinction memory retrieval.

## Silencing IL-to-BLA projection neurons impairs context-dependent extinction memories

To establish the IL projection system necessary for extinction memory, we used the PSAM-PSEM chemogenetic system^18^ to manipulate action potential output. We first made whole-cell patch-clamp recordings of PSAM-GlyR expressing neurons in anesthetized mice (n=4 cells from 4 mice), which revealed intraperitoneal administration of the ligand 817 (1 mg/kg) rapidly reduced spike rates and the membrane potential variance in PSAM GlyR-expressing IL pyramidal neurons (**Figure 2A-B**). Results using fiber photometry Ca^2+^ imaging in awake mice also found that the frequency and amplitude of Ca^2+^ transients in IL excitatory neurons were decreased with 1 mg/kg and 0.3 mg/kg of 817. Administration of 817 alone, in the absence of the IL expression of the PSAM-GlyR receptor, produced no change in Ca^2+^ transient frequency or amplitude (**Extended Data Figure 4A-C**).

**Figure 2.**
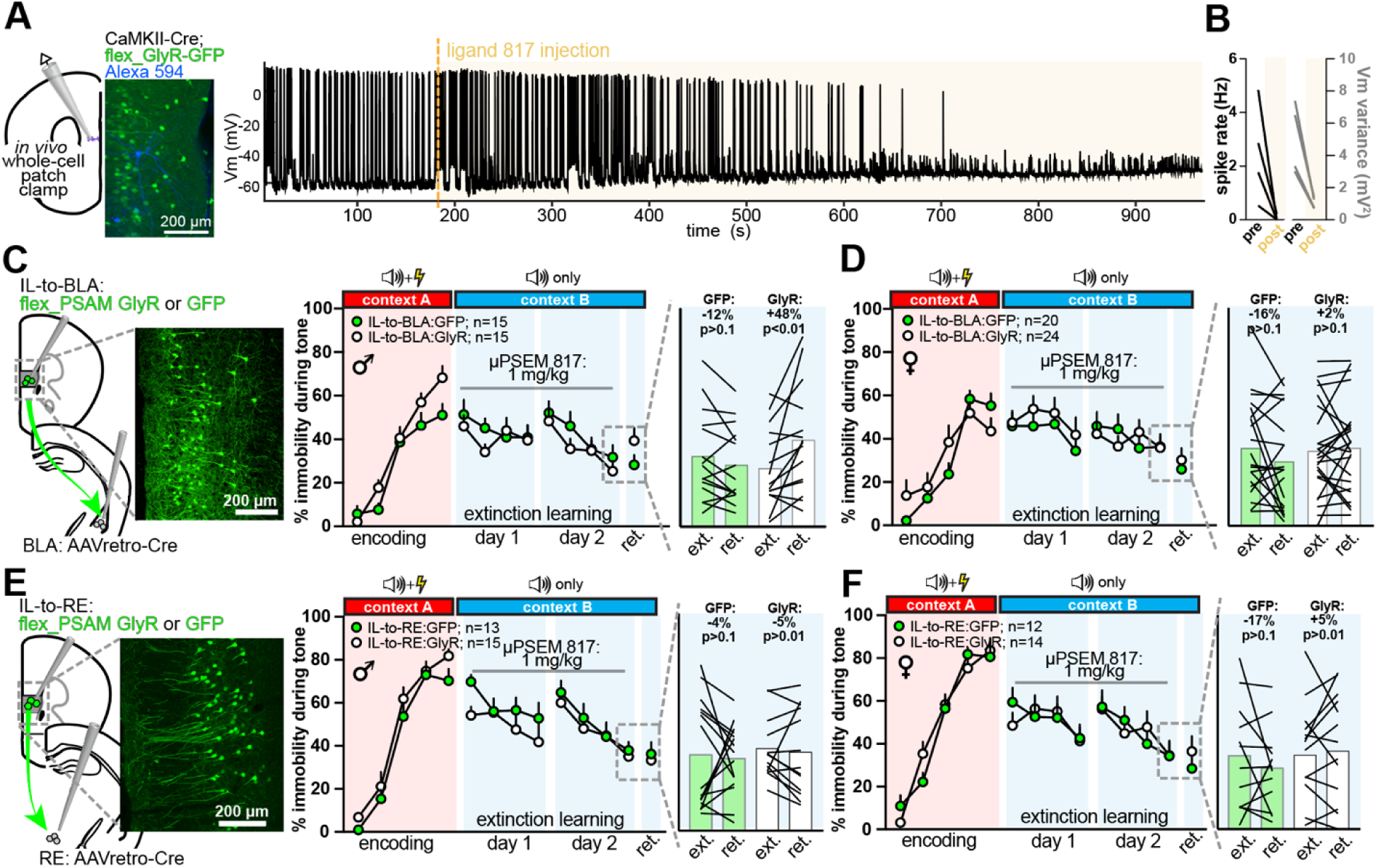
Chemogenetic neuronal silencing reveals activity within IL-to-BLA neurons is required for extinction memory in male but not female mice. **A**) Whole-cell patch clamp recordings of PSAM-expressing IL neurons (left) made from anesthetized mice show administration of ligand 817 (1 mg/kg, i.p.) rapidly and robustly silences action potential output (post-injection recording shaded orange). **B**) Somatic spike rate and subthreshold membrane (Vm) variance were reduced within minutes of 817 injection (n=4 cells from 4 mice; one-tailed Student’s paired t-test, p=0.03 for spike rate; p=0.01 for Vm variance). **C**) Mice were injected with AAVretro-Cre in the BLA and Cre-dependent rAAVs expressing PSAM-GlyR or GFP into IL. After recovery, 817 (1 mg/kg, i.p.) was administered 30 min prior to both extinction learning sessions. IL-to-BLA silencing left encoding and within-session extinction intact (two-way ANOVAs, P>0.05 for post-hoc tests) but reduced extinction memory retrieval in male mice (n=15 mice/group, two-tailed Student’s paired t-test, p<0.01 for GlyR mice). **D**) The same experiment in female mice (n=20 GFP mice and n=24 GlyR mice) produced no significant change during extinction learning (two-way ANOVAs, P>0.05 for post-hoc tests) or at extinction memory retrieval (two-tailed Student’s paired t-test, p>0.05 for each group). **E-F**) Inhibition of IL-to-RE neurons during extinction learning produced no change in extinction memory retrieval in male mice (**E**; n= 13 GFP mice and n=15 GlyR mice; p>0.05 for all comparisons) or in female mice (**F**; n=12 GFP mice and n=14 GlyR mice; p>0.05 for all comparisons).

IL projections to the basolateral amygdala (BLA) and nucleus reuniens of the thalamus (RE) have each been implicated in extinction learning and extinction memory retrieval^13,16^. These two populations are almost entirely non-overlapping in IL, with IL-to-BLA neurons residing in the superficial layers and IL-to-RE neurons residing in the deeper layers^19^. To restrict expression of PSAM-GlyR chemogenetic receptors to one projection population, we injected AAVretro-Cre^20^ bilaterally into the target region followed by injection of AAV2/1-Syn-flex-*rev*-EGFP or AAV2/1-Syn-flex-*rev*-GlyR-2a-EGFP bilaterally into IL. Ligand 817 (1 mg/kg, i.p.) was administered 30 minutes prior to both extinction learning sessions to all mice. In male mice (n=15-16 mice/group), inhibition of IL-to-BLA neuronal activity during extinction learning sessions impaired subsequent extinction memory retrieval. Specifically, GlyR-expressing mice showed 48% increase in their immobility during the tone at extinction memory retrieval as compared to the last trial block from extinction learning session 2 (**Figure 2C**, p<0.01, paired t-test). The effect of silencing IL-to-BLA neurons on extinction memory retrieval was absent in female mice (**Figure 2D**, n=20-22 mice/group, p>0.5 for all comparisons). We used the same approach to silence IL-to-RE neurons in male (n=13-15 mice/group) and female (n=12-14 mice/group) mice during extinction learning and found no change in extinction memory retrieval in mice of either sex (**Figure 2E-F**). Collectively, these data show that neural activity in IL-to-BLA neurons is required during extinction learning in male but not female mice for later extinction memory retrieval.

## Learning-related synaptic formation and clustering on IL-to-BLA projection neurons

Somatic spiking activity is a critical intracellular signal for both Hebbian^21^ and non-Hebbian^22,23^ forms of synaptic plasticity. Such plasticity requires both RNA and protein synthesis, and IL infusion of inhibitors to these processes during extinction learning impairs extinction memory retrieval^24,25^. To establish the presence and magnitude of synaptic plasticity generated on IL neurons during extinction learning, we used a dual-recombinase AAVretro strategy (**Figure 3A, Extended Data Figure 5A**) to reconstruct synaptic densities on perisomatic branches from IL-to-BLA and IL-to-RE projection neurons. As spine head size correlates strongly with synapse weight^26,27^, spine neck geometry controls spine compartmentalization from the parent dendrite^28^ and both features contribute to long-term synaptic potentiation^29,30^, we reconstructed the spine head volumes and neck lengths of individual spines on these branches.

**Figure 3.**
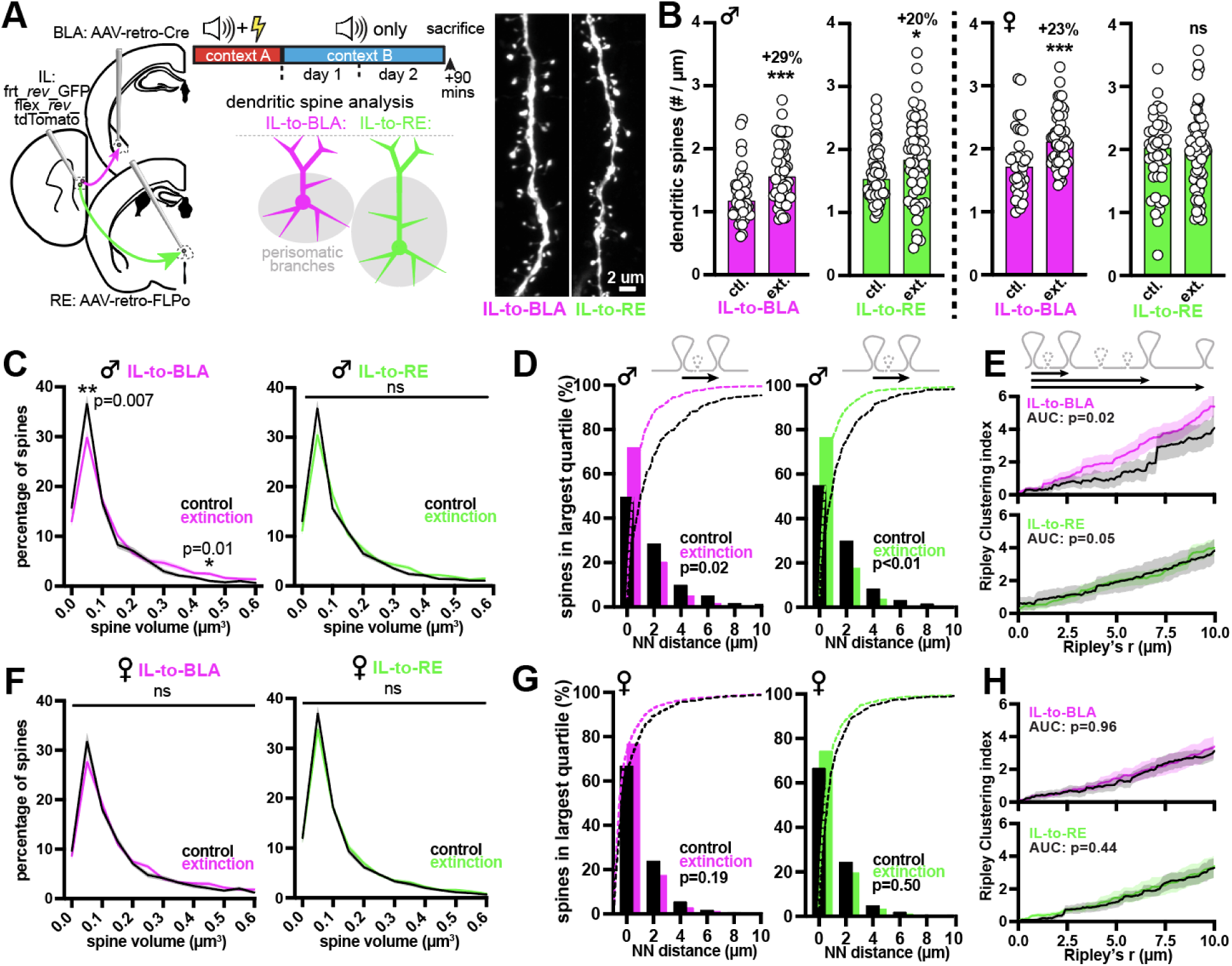
Extinction learning is associated with new synapse formation, synaptic structural plasticity, and clustering of large synapses in male but not female mice. **A**) Dual AAVretro strategies were used to drive tdTomato in IL-to-BLA neurons and GFP in IL-to-RE neurons; mice were sacrificed 90 min. after the second extinction session and dendritic segments were analyzed. **B**) Spine densities were increased on perisomatic branches of IL-to-BLA (45 control branches from 4 mice; 50 extinction branches from 5 mice: two-tailed Mann Whitney test, p=0.0002) and on IL-to-RE neurons (60 control branches from 5 mice; 59 extinction branches from 5 mice: two-tailed Mann Whitney test, p=0.023) from male mice (left plots). Similar density increases were evident on IL-to-BLA neurons (38 control branches from 4 mice; 53 extinction branches from 5 mice: two-tailed Mann Whitney test, p<0.0001) but not IL-to-RE neurons from female mice (right plots; IL-to-RE: n=41 control branches from 4 mice; n=68 extinction branches from 5 mice: two-tailed Mann Whitney test, p=0.88). **C**) Spine volume showed a shift away from small spine volumes and towards larger spine volumes on IL-to-BLA branches (two-way ANOVA, 0.05 bin, p=0.007; 0.45 bin, p=0.01) but not IL-to-RE neurons from male mice (two-way ANOVA, p>0.05 for all comparisons). **D**) Spines from male mice in the largest quartile had shorter nearest neighbor distances on branches from IL-to-BLA and IL-to-RE neurons (Fisher’s Exact test: IL-to-BLA, p=0.02; IL-to-RE, p<0.01). **E**) 1-D Ripley’s analyses showed enhanced clustering of groups of large spines on IL-to-BLA (K-S test of individual branch cluster scores, n=33 control branches and n=47 extinction branches, p=0.02) and moderately on IL-to-RE neurons (K-S test of individual branch cluster scores, n=56 control branches and n=58 extinction branches, p=0.05). **F-H**) The same analyses of spine head volume distributions, nearest neighbor distances, or intrabranch clustering revealed no significant effects of extinction learning in female mice (p>0.05 for all comparisons).

In male and female mice, a single extinction learning session produced no change in dendritic spine densities in either projection class (**Extended Data Figure 5B**) and no change in spine morphologies on IL-to-BLA neurons but induced a modest shift toward larger spine head volumes on IL-to-RE neurons from male mice (**Extended Data Figures 5 and 6**). However, the second extinction learning session produced robust elevations of dendritic spine densities (∼30%) on the perisomatic dendrites of IL-to-BLA neurons from male and female mice (**Figure 3B**: control vs. extinction, p<0.001 for both sexes), and on IL-to-RE neurons selectively from male mice **(Figure 3B**: control vs. extinction, p=0.0.2 for males, control vs. extinction, p=0.88 for females). The increase in spine densities was accompanied by a shift towards larger spine head volumes and neck lengths on IL-to-BLA but not IL-to-RE neurons from male mice (**Figure 3C).** Spine head and neck morphology was effectively unchanged on either projection class in female mice (**Figure 3F, Extended Data Figure 6B**). Thus, the second extinction learning session, which is required for persistent context-dependent extinction memory retrieval, is associated with synaptic rewiring and structural plasticity consistent with strengthening of synapse weights on IL-to-BLA neurons in male but not female mice.

Past work has suggested that clustering of synaptic inputs promotes distinct forms of dendritic integration compared to spatially distributed synapse distributions^31^. To examine whether the learning-related changes in spine density and spine head size were associated with increased spatial clustering of large synapses, we performed a nearest neighbor proximity analysis of synapses from the largest quartile along branch segments from IL-to-BLA and IL-to-RE neurons. Nearest neighbor intrabranch distances were decreased on both IL-to-BLA and IL-to-RE branches from male but not female mice (Fisher’s exact test; p<0.05 for both projection classes in male mice; p>0.1 for both projection classes from female mice; **Figure 3D,G**). Increased clustering of large spines on IL-to-BLA and to a lesser extent on IL-to-RE branches from male but not female mice was found using a 1-D Ripley’s analysis, which allows analysis along a continuum of distances and examines neighborhoods of large spines rather than simply pairs (**Figure 3E-H**).

Last, we asked whether the altered propensity for spine plasticity across male and female mice during extinction learning might arise from differences in the wiring of IL afferent axons. Extinction-related signals arise early during learning in BLA neurons in neurons that project to IL^32^, so we expressed AAV2/1-CamKIIa-Cre and AAV2/1-Syn-flex-*rev*-mGFP-2a-Synaptophysin_mRuby^33^ in BLA excitatory neurons and analyzed the density of putative synapse boutons along BLA axons in layer III of IL (**Extended Data Figure 7A-B**; see **Methods**). We found that male and female mice have effectively identical bouton densities on BLA axons in IL (ANOVA, p>0.05 for all comparisons; **Extended Data Figure 7C-D**), arguing against coarse sex differences in afferent connectivity as a factor shaping reduced plasticity on IL neurons in female mice. Collectively, these results show the second extinction learning session reorganizes synaptic connectivity in terms of synapse density, synapse size, and intrabranch spatial clustering on branches from male but not female mice.

## GRIN2B in IL-to-BLA neurons controls extinction memory retrieval and learning-related synapse plasticity

The formation of new synapses and the structural synaptic plasticity of existing spines is governed by NMDA receptors^34,35^. The NMDAR subunit GRIN2B influences synaptic calcium entry and channel kinetics^36^ and supports synapse plasticity by its unique C-terminal domain, which binds CaMKIIa^37^ – a critical intracellular structural trigger for synapse plasticity^38,39^. Using postembed immunoelectron microscopy, we found GRIN2B localizes in IL synapses to the postsynaptic density, and its expression in IL did not differ between male and female mice (**Extended Data Figure 8A**).To establish a functional role for GRIN2B, we paired our projection-specific viral recombinase strategies with CRISPR-generated mice containing loxP sites flanking exon 4 of the *grin2b* gene (hereafter *grin2b^fl/fl^*). By injecting AAVretro-FlpO into the BLA and AAV2/1-EIFa-frt-*rev*-Cre into the IL of *grin2b^fl/fl^* mice, the deletion of *grin2b* can be achieved on demand over a relatively short timescale in adult mice and limited to a highly specific subset of cortical neurons. We observed no leak from our viral frt-*rev*-Cre approach in the absence of AAVretro-FlpO (**Extended Data Figure 8B**), and validated *grin2b* gene deletion with fluorescent *in situ* hybridization using probes against *flpo* and *grin2b* and with anti-Cre immunofluorescence in IL slices. Cre-containing IL projection neurons show ∼80% reduction in *grin2b* mRNA relative to FlpO+/Cre– IL-to-BLA and IL-to-RE neurons (**Figure 4A-B**).

**Figure 4.**
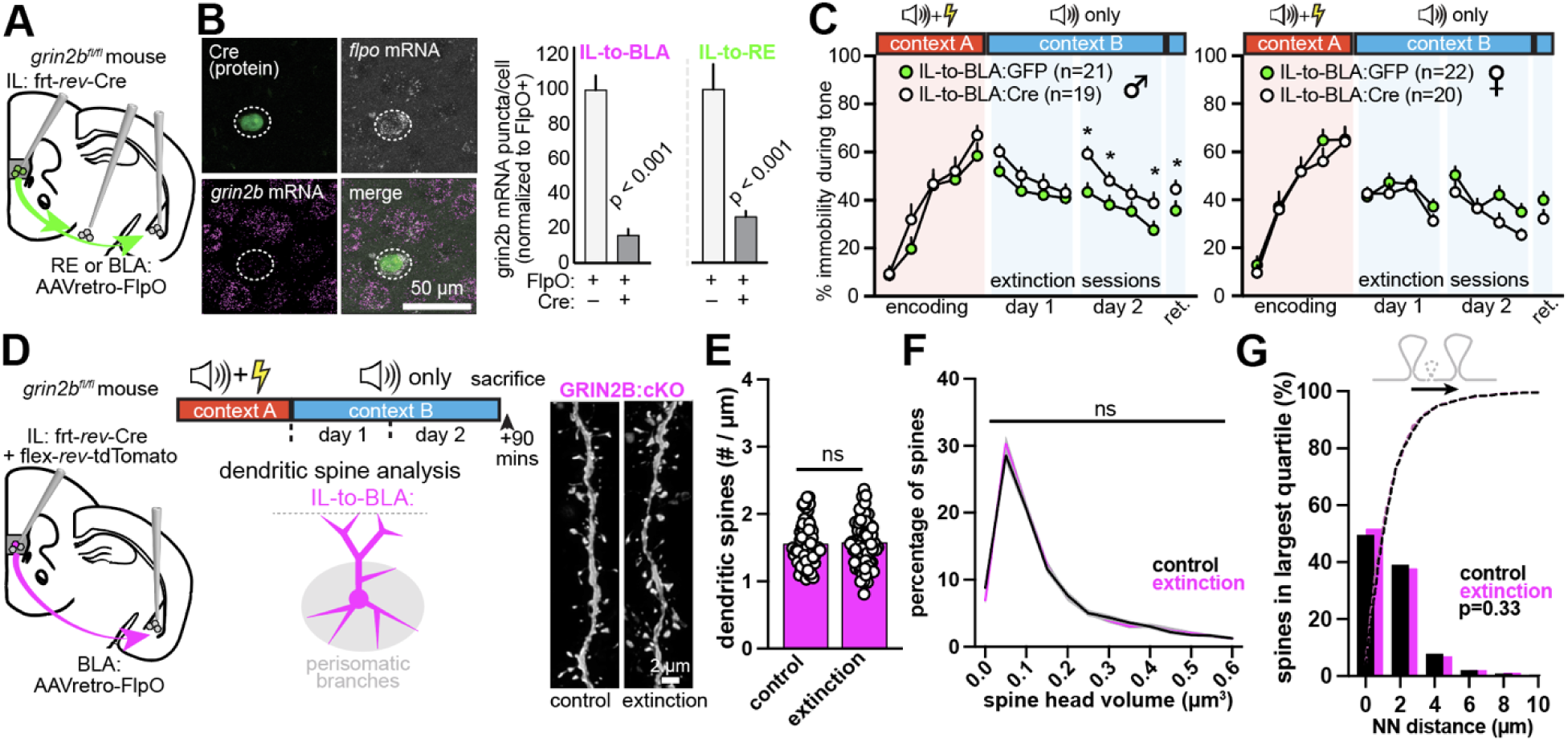
GRIN2B in IL-to-BLA neurons mediates extinction memory in male but not female mice. **A**) Pairing a dual recombinase viral strategy with a *grin2b^fl/fl^* mouse permitted the deletion of *grin2b* solely from IL-to-BLA neurons. AAVretro-FlpO was injected into the BLA, and FlpO-dependent Cre was injected into IL. **B**) Efficiency of gene deletion was assessed using fluorescent in situ hybridization with probes for *flpo* and *grin2b* and immunofluorescent labeling of Cre protein. In both IL-to-BLA and IL-to-RE projection neurons, *grin2b* mRNA in *flpo*+/Cre+ was reduced ∼80% relative to *flpo*+/Cre– neurons (n=30 *flpo*+/Cre+ cells and n=30 *flpo*+/Cre-cells taken from 2 IL-to-BLA mice; n=81 *flpo*+/Cre+ cells and n=32 *flpo*+/Cre– cells taken from 2 IL-to-RE mice; t-test, p<0.0001 for each projection class). **C**) Deletion of *grin2b* in IL-to-BLA neurons left behavior during encoding and the first extinction session unchanged, but increased immobility to the tone during the second extinction session and at extinction memory retrieval (left: Ext 2; two-way ANOVA, p=0.003 for block 1 and p=0.02 for block 2; extinction memory retrieval, unpaired t-test, p=0.03; n=21 GFP mice and n=19 Cre mice). No changes were evident with the same manipulation in female mice (right; p>0.05 for all comparisons; n=22 GFP mice and n=20 Cre mice). **D**) IL-to-BLA neurons were labeled and GRIN2B deleted using the same dual-recombinase strategy as in **A**, mice were sacrificed after extinction session 2, and branche from male mice exposed to control (no shock; n=67 branches from 5 mice) or extinction conditions (n=74 branches from 6 mice) were reconstructed. **E-G**) With GRIN2B deletion in IL-to-BLA neurons, dendritic spine density was unchanged (**E**; p=0.51, unpaired two-tailed t-test), spine head volume distributions were unchanged (**F**; n=67 branches from 5 control mice, n=74 branches from 6 extinction mice, two-way ANOVA, p>0.1 for all comparisons), and clustering of large spines was absent (**G**; Fisher’s exact test, p=0.33).

We next asked if deletion of *grin2b* impaired extinction memory retrieval. Bilateral deletion of *grin2b* from IL-to-BLA neurons did not alter tone-dependent immobility rates during encoding or during the first extinction learning session. However, *grin2b* deletion from IL-to-BLA neurons increased immobility rates to the tone during the second extinction learning session relative to GFP-expressing mice (trial blocks 1 and 2, p<0.05 for both; **Figure 4C**). The effect of *grin2b* deletion persisted to the extinction memory retrieval phase (p<0.05), where *grin2b*-deleted mice showed higher rates of immobility to the tone than *grin2b*-intact mice. By contrast, *grin2b* deletion from IL-to-BLA neurons in female mice did not alter behavior during encoding, across either extinction learning session or at extinction memory retrieval (p>0.1 for all comparisons). Consistent with the negative results from chemogenetic silencing experiments, deletion of *grin2b* from IL-to-RE neurons in male or female mice produced no change of tone-associated immobility in any task phase (**Extended Data Figure 8C**). Collectively, these results show that GRIN2B is necessary in IL-to-BLA neurons for extinction learning and extinction memory retrieval.

To establish if GRIN2B impaired extinction concomitant with blocking learning-related synapse plasticity, we used the same dual-recombinase approach described above to delete *grin2b* strictly from IL-to-BLA neurons and reconstructed dendritic spines after the second extinction learning session in male mice. IL-to-BLA deletion of *grin2b* completely blocked the increase in spine density seen after the second extinction learning session (**Figure 4D**), the learning-related shift toward larger spine volumes (**Figure 4E**), and the clustering of large inputs along dendritic branches (**Figure 4F**). Spine densities, spine head morphologies, and spine clustering from GRIN2B-deleted mice were effectively identical to those from control B6 mice (**Extended Data Figure 8D-G**), strengthening the evidence for coordinate GRIN2B-dependent control over learning-related plasticity and memory retrieval in male but not female mice.

## DISCUSSION

Here we used *in vivo* Ca^2+^ imaging, neuronal activity silencing, synapse reconstructions, and projection-specific gene deletions to show a critical role for synaptic plasticity on IL-to-BLA projection neurons during context-dependent extinction learning and extinction memory retrieval. Our results show that multiple aspects of a synapse plasticity mechanism necessary for extinction memory in male mice are absent in female mice. These include the requirement for activity in IL-to-BLA projection neurons, learning-related spine structural plasticity and clustering, and signaling downstream of the NMDA receptor subunit GRIN2B within IL-to-BLA neurons. Although both male and female mice showed similar patterns of tone-dependent neural activity responses and within– and across-session neural activity dynamics in IL, the mechanisms recruiting IL neurons during extinction learning appear distinct between male and female mice. Because mice from both sexes had nearly identical behavioral responses to extinction learning and at extinction memory retrieval, these findings imply different synaptic and cellular mechanisms underlie the same behavioral function across the sexes.

Our recordings show IL contains tone-responsive excitatory neurons during each task phase, and that individual neurons displaying tone representations leave and join the active ensemble at high rates within and across sessions. These results are broadly consistent with previous reports of dynamic neural codes underlying performance in cortical and hippocampal circuits^14,15^. Neurons responses to the conditioning tone in early portions of a learning session are inhibited at later points and vice versa, suggesting the IL tone ensemble may be controlled by dynamic routing of excitation or inhibition to individual neurons during learning and memory retrieval. Across-session dynamics were reduced after the second session, with an increase in tone-responsive neuron participation from the second extinction learning session to extinction memory retrieval seen in each animal. That such a change in dynamics occurs alongside the induction of synaptic plasticity in IL strongly implies the phenomena are related.

Our chemogenetic silencing experiments show that projections to the amygdala are critical during extinction learning for later extinction memory retrieval. Such a result is consistent with past work that has manipulated IL-to-BLA neurons using different strategies^16,17,40^, none of which examined the relevance of this pathway for behavior in female mice. That this pathway is indispensable for male but not for female mice is consistent with the synapse structural plasticity and increased clustering of large synapses seen selectively on IL-to-BLA projection neurons from male mice.

Functionally, the increased clustering of large synapses may result from increased local dendritic spine crosstalk^41^and should alter the integration of inputs on these branches by the facilitation of dendritic spike generation^31^. GRIN2B has been shown to control synaptogenesis and synaptic plasticity though its C-terminal tail^34,42^, which is unique among NMDAR subunits in its ability to bind CaMKIIa^39,43,44^. Recent work has shown the GRIN2B-CamKIIa interaction serves as an intraspine structural trigger for synaptic potentiation^38^. Our projection-specific deletion results are consistent with a critical role for GRIN2B in IL-to-BLA neurons mediating spine plasticity and extinction memory formation in male mice. The lack of a behavioral effect after deleting *grin2b* in these neurons from female mice is consistent with our negative IL-to-BLA silencing results and with the diminished spine structural plasticity and clustering seen on these neurons in female mice.

Although IL has long been implicated as a controller of extinction memories^4–6^, the cellular, synaptic, and molecular logic by which specific IL neurons are recruited during learning have remained unresolved. That the GRIN2B-dependent synaptic plasticity mechanism used by IL-to-BLA neurons to form and store extinction memories in male mice is absent in female mice suggests that extinction learning is driven by distinct mechanisms across the sexes. The disparate nature of these mechanisms may be conserved across species, as functional magnetic resonance imaging of men and women show differential activation of frontal cortical networks during fear learning and during extinction^45^.

Future work should expand the behavioral repertoire by which extinction memories may be expressed across male and female mice^46^ and explore the circuit, cellular, and molecular mechanisms that control extinction memory expression in females, including the potential modulation by circulating hormones^47^. Although estrogen is well-known to modulate structural plasticity in cortical circuits^48–51^, past work has shown local brain estrogen levels may be comparable or higher in males^52,53^. Moreover, estrogen-induced plasticity in cortical neurons depends on NMDA receptors^54^, which should heighten, rather than dampen, the dependence of NMDARs on plasticity and memory formation in female mice. Independent of endocrine status, our results have broad implications for conceptual views on the mechanisms of information processing in cortical circuits. They also emphasize the importance of developing precision medicine approaches for treating mental health disorders as opposed to one-size-fits-all strategies, including recent clinical trials using GRIN2B subunit-specific modulators (e.g., NYX-783) for the treatment of post-traumatic stress disorder.

## Supporting information

Supplemental Statistical Tables

## ONLINE METHODS

### Animals

Adult mice (3-6 mos. of age) of both sexes were used. Lines used in this study include B6 mice (C57BL6/J; JAX Strain # 000664), Ai75d mice (B6.Cg-Gt(ROSA)26Sor^tm75.1(CAG-tdTomato*)Hze^/J; JAX Strain # 025106), and *grin2b*^fl/fl^ mice (C57BL/6J-^Grin2bem5Lutzy^/J; JAX Strain # 032664), all of which are kept on the C57BL6/J background and are commercially available from The Jackson Laboratory. Mice were kept in polysulfone cages (30.8 cm x 30.8 cm x 16.2) with a center partition at 20-22.2°C and 40-50% humidity with food and water available ad libitum.

### Intracranial injections and GRIN lens implantation

Recombinant adenoassociated viral vectors (rAAV; serotype 2/1 or AAVretro), obtained from Addgene, University of Pennsylvania, Janelia Research Campus, or UNC-Neurotools, were used to drive expression of calcium indicators, recombinases, fluorescent reporters, or chemogenetic receptors. Viral specificity was tested in wild-type mice to confirm Cre or FlpO recombinase dependency. Viral constructs (20-200nL) were injected as previously described^19,55,56^ using glass pipettes (∼30 µm diameter tips) with Drummond Nanoinjectors driven by micromanipulators (Sutter) on mice under isoflurane anesthesia (1-3%, mixed with oxygen) positioned in a Kopf stereotactic apparatus.

For experiments in which mice were implanted with GRIN lenses (Inscopix Proview DC Integrated Lens 1 mm diameter x 4 mm length), viral injections and lens implantations were performed in the same surgery. The integrated lens and baseplate were affixed to the skull with C&B-Metabond Quick Adhesive Cement System (Parkell). Following surgery, mice were administered carprofen (5 mg/kg, 0.1 ml/10 g of body weight), Ethiqa (1.3 mg/ml, 0.01 ml/ 10 g body weight) and ropivacaine (1 mg/ml; topical) to facilitate pain management and recovery from surgery and were allowed to recover in a heated cage prior to returning to their home cage. All mice were allowed at least three weeks of recovery prior to behavioral testing.

### Context-dependent Fear Extinction

Mice were handled by an experimenter for 2 minutes for at least three consecutive days prior to the extinction paradigm. For calcium imaging experiments, mice were habituated to the microscope and tether for 30 min each of the 3 days. All behavioral tests were performed in the mouse’s light cycle (approximately 2 hours after lights on), and mice assigned to different group conditions were run in interleaved batches to control for within session timing. All experimenters were blind to the condition during the behavioral sessions.

Encoding and extinction was performed in Lafayette Instruments mouse boxes (interior dimensions: 45.75 x 45.75 x 55.1 cm) equipped with cameras (3.75 frames/sec) and analyzed with FreezeFrame software. For each session, mice were allowed 2 mins of contextual investigation prior to the onset of any tones then 4kHz pure tones were played at 80dB for 20 seconds each. For encoding (Context A), 3 tones were played to determine baseline response to the tone-only followed by 5 tones paired with a 1 second, 0.8mA foot shock. For extinction sessions (Context B), a series of 12 randomly dispersed tones were played across a 20-minute session. For retrieval and renewal (Context B), 4 evenly dispersed tones were played during a 6-minute session. In the experiments shown in Figure S1, mice were placed back into the chambers for retrieval and renewal tests exactly one week after the original tests. The differences between the contexts included odor, light, and floors/walls: (1) Context A = circular, striped wall inserts, almond extract, all lights on, and cleaned with 70% ethanol solution between animals; (2) Context B = opaque white Plexiglas floor insert with attached Velcro pieces, banana extract, all lights dimmed and cleaned with Virkon solution between animals.

### *In vivo* Ca^2+^ imaging and analysis

Neuronal activity was measured using GCaMP6f signals as a proxy for spiking activity and recorded by GRIN lenses implanted 2.5 mm deep at a 15-degree angle, just dorsal to IL. For each session, optical recordings (20 Hz; 4.0-4.8 gain; 0.4-0.8mW LED power; collected from Inscopix nVue2.0 IDAS) were transformed using the Inscopix IDPS software. Specifically, imaging frames were downsampled to 10HZ and subject to spatial and temporal downsampling, spatial bandpass filtering and motion correction before cell identification of putative neurons using the constrained non-negative matrix factorization (CNMFe) algorithm^57^. The traces of all cells were manually inspected to exclude false-positive or false-negative cell mask allocation. The identified cells from CNMFe were co-registered across the behavioral sessions using the longitudinal registration with a minimum correlation of 0.50. Raw calcium traces were obtained by averaging all the pixel values in each mask. Each raw trace file was then separated by timestamps for extinction day 1, extinction day 2 and retrieval and aligned with their respective behavior files for tone epochs and freezing behavior to the nearest timestamp. Each cell was then individually z-score normalized to the entire behavior session.

### Event-related neuronal activity

Calcium activity data were analyzed using the Python programming language (https://www.python.org/) (code available on request). To compute event-related calcium responses to presentation of the tone over the course of each extinction session and retrieval, the 12 tone epochs were blocked by early (first 3 tones for extinction session and first tone for retrieval), all tones, or late (last 3 tones for extinction sessions and last tone for retrieval). Neurons exhibiting a significant event related change (z-score change from tone onset greater than 0.5 within 3 seconds post-tone onset for a minimum of 10 consecutive frames) for early, all, or late tones were expressed as the percentage of all neurons recorded for that session. The percentage overlap between sets of event-responsive neurons across sessions was calculated as the proportion of co-registered neurons.

### In vivo patch clamp electrophysiology and fiber photometry

To validate effective neuronal silencing by the PSAM-PSEM system, patch clamp recordings were made from head-fixed, anesthetized mice. Recordings were made exactly as in^58^, except for the use of anesthesia (1-2% isoflurane mixed with oxygen) and the attachment of thin polyethylene tubing connected to a syringe containing the chemogenetic ligand 817 (given at 1 mg/kg) to the intraperitoneal cavity though a 27-gauge needle. After somatic break in, we waited approximately 2 minutes prior to administration of 817 to establish baseline values for somatic spiking activity and membrane potential variance. Whole-cell recordings were made throughout the session with no holding current. Recorded neurons were identified as expressing PSAM-GlyR by post-hoc visualization of biotinylated Alexa 594 (as shown in **Fig. 2**).

Fiber photometry recordings were made from awake mice co-expressing PSAM-GlyR-GFP and jRCaMP in IL excitatory neurons. Ca^2+^ transients were analyzed using a time-division multiplexing strategy as in^59^ with the GlyR-GFP signal as a static movement control channel. Recordings were made in sessions just prior to and immediately after i.p. injection of ligand 817 (1 mg/kg or 0.3 mg/kg). Spontaneous peaks were detected in the movement-corrected jRCaMP ΔF/F signal using Matlab function findpeaks(), with peak prominence set to the median absolute deviation of the control session prior to ligand injection. Analyzing Ca^2+^ transient detection using a range of other peak prominence values yielded qualitatively similar results.

### Tissue collection, immunolabeling/in situ hybridization, microscopy, and analysis

Mice were anesthetized with Tribromoethanol (800 mg/kg, i.p.) and transcardially perfused with ice-cold 4% depolymerized paraformaldehyde (EMS) in 0.1 M PB. For immunofluorescent protein labeling, coronal slices were cut on a Vibratome (50-60 µm-thick) and were incubated in blocking buffer containing 2% BSA and 0.3% Triton X-100 followed by incubation in the relevant primary antibodies (Guinea Pig anti-cFos, Synaptic Systems; Guinea Pig anti-Cre, Synaptic Systems; Rabbit anti-Grin2b, Sigma) at 4°C for 24-48 hours. Slices were washed in buffer containing 0.3% Triton X-100 and followed by incubation of the fluorescent-conjugated secondary antibodies diluted in the same blocking buffer. For immunofluorescent imaging of spines, GFP and tdTomato signals were not enhanced with antibodies. Slices were mounted onto microscope slides and coverslipped with Vectashield.

For postembed immunogold labeling, mice were perfused with 4% paraformaldehyde + 0.125% glutaraldehyde and tissue punches (3 mm diameter) were rapidly vitrified with a Leica ICE high-pressure freezer. Tissue blocks were freeze-substituted, embedded into HM20 resin, polymerized by UV light, cut (70 nm-thick) on a ultramicrotome, and collected onto carbon-coated Formvar slot Grids (EMS). Postembed immunolabeling was performed as previously described^60^ with prior etching in a saturated solution of NaOH in absolute ethanol for 2–3 s followed by rinses in distilled water^61^ prior to the conventional immunogold labeling protocol. Grids were imaged with a JEOL 1230 transmission electron microscope and images were collected with a CMOS camera (Hamamatsu).

RNAscope was performed as described previously^55^ under RNAse-free conditions/solutions using cryostat coronal sections (14 µm-thick) and probes to GRIN2B and FlpO (ACDBio). Following probe hybridization, sections were processed for Cre immunofluorescence using a polyclonal antibody (Synaptic Systems, described above) followed by an Alexa 488-conjugated secondary.

Confocal microscopy for dendritic and axonal analysis was performed using a Leica Confocal Microscope (SP8 or Stellaris) equipped with multiple objectives, laser lines, and PMT/HyD photodetectors. Dendritic branches and spines were imaged with a 63x 1.4 NA objective, at 0.05 x 0.05 x 0.1 µm voxel sizes, and processed through Lightning Deconvolution. Dendritic spines and axons were analyzed in NeuronStudio^62^, which allows a semiautomated analysis of branch/cable and characteristics individual spine head and neck morphologies. For the reconstruction of axons, putative boutons were identified as nodes with diameters >1.5 standard deviation from the cable mean. Fluorescent *in situ* hybridization signals were captured with a 63x 1.4 NA objective at 0.18 x 0.18 x 0.3 µm voxel sizes. Individual *grin2b* puncta were segmented in Fiji and counted within a 20µm-diameter circular mask centered over the nucleus. Each cell was characterized as FlpO± and Cre± based on the presence or absence of the relevant fluorescent signal, and counts were normalized to control means from intact neurons belonging to the same projection class (i.e., FlpO+/Cre-) to estimate the magnitude of gene expression knockdown.

### Statistical Analyses

Behavior analyses were performed blind to the experimental conditions using automated Actimetrics software to determine immobility epochs defined by a minimum absence of movement for 1s, with the threshold adjusted on a case-by-case basis. Immobility during the tone during encoding and extinction task phases was analyzed separately with mixed-model two-way repeated measures ANOVAs with treatment and trial blocks as factors and Holm-Sidak posthoc tests performed with corrections for multiple comparisons. Retrieval sessions were analyzed using unpaired t-tests. In experiments for which we have a strong directional hypothesis (e.g., 2 extinction sessions vs. 1 extinction session, or somatic spike rates in silencing experiments), one-tailed t-tests were used; in experiments where we have no directional hypothesis (e.g., IL-to-BLA *grin2b* deletion), two-tailed t-tests were used. Dendritic spine densities were analyzed separately for each sex, extinction learning session, and projection class using parametric or non-parametric two-tailed t-tests (depending on normality test results). Dendritic spine head volumes and neck lengths were analyzed with mixed-model two-way repeated measures ANOVAs with condition and bin size as main factors and Holm-Sidak posthoc tests were performed with corrections for multiple comparisons. For clustering analyses, branch segments were excluded if they were shorter than 20 µm to minimize edge artifacts. Nearest neighbor distributions were binned (2 µm) and analyzed with a Fisher’s Exact test, and intrabranch clustering was analyzed using a Ripley’s 1D K test where:

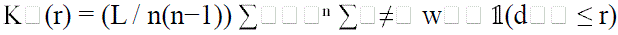

Where n is the number of objects, L is the total dendritic segment length, d□□ is the distance along the dendrite between synapse i and j, r is a radius, and w□□ is an edge correction. To compare the output to a Poisson process, the L-function was implemented as: L(r) = K□(r) / 2 and plotted as H(r) = L(r) – r. For every individual cable, we calculated the Ripley’s L-function curve across a 0-10 µm range (inspired by^41,63^). To determine if the groups differed, we calculated the area under the curve (AUC) for each specific segment, and tested differences across AUC distributions using a nonparametric Kolmogorov-Smirnov test.

## DATA AND CODE AVAILABILITY

All data and code that supports this work are available at Figshare (*live doi to be inserted and made available at publication*). Requests for additional information can be made to

## ACKNOWLEDGEMENTS

We thank Drs. R. Burgess, G. Howell, M. Joy, K. O’Connell, and P. West for helpful discussion and comments; Drs. Fabio Cacciapaglia and Sinem Erisken from Bruker/Inscopix, Dr. Jacqui White from the Center for Biometric Analysis at The Jackson Laboratory, and Dr. Philipp Henrich from Light Microscopy Core at The Jackson Laboratory. We thank E. Nickerson and L. Lepeak for assisting with experiments and for critical feedback.

## AUTHOR CONTRIBUTIONS

Conceptualization, K.G and E.B.B.; Methodology, K.G, J.P.T, G.O, L.K.W, A.D., L.C, X.Z, and E.B.B; Formal Analysis, K.G, J.P.T, G.O, L.K.W, A.D., L.C, X.Z, and E.B.B; Investigation, K.G, J.P.T, G.O, L.K.W, A.D.,L.C, X.Z, and E.B.B; Writing – Original Draft, K.G. and E.B.B; Writing – Reviewing & Editing, K.G, J.P.T, G.O, L.K.W, A.D., L.C, X.Z, and E.B.B; Supervision, E.B.B; Funding Acquisition, E.B.B.

## EXTENDED DATA FIGURES

**Extended Figure 1.**
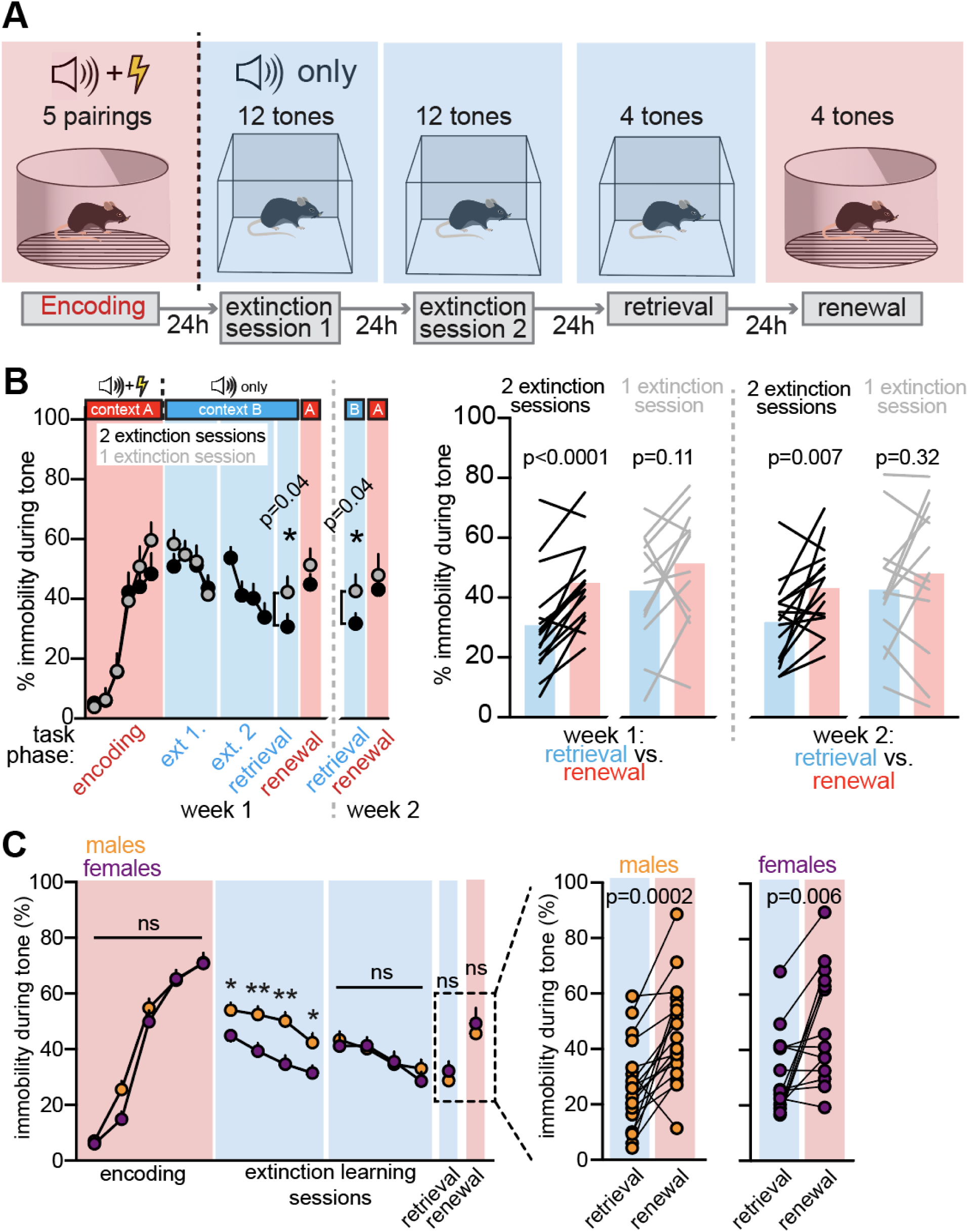
Two sessions of extinction learning are required for persistent context-dependent extinction in male and female mice. **A**) Schematic representing the context-dependent extinction paradigm. **B**) Mice required two extinction learning sessions to maintain persistent, weeks-long context-dependent discrimination. Mice were subjected to a single extinction learning session (grey symbols, n=13) or to two extinction learning sessions (black symbols, n=16 mice). Across-group comparisons show weaker extinction performance at an extinction memory probe trial (p=0.04, one session vs. two sessions), and on a second probe trial one week later (p=0.04, one session vs. two sessions). Discrimination between the encoding and extinction contexts was significant in mice exposed to two extinction sessions (week one: p<0.0001; week two: p=0.007; paired t-tests) but not at either time point for mice exposed to a single extinction learning session (week one: p=0.11; week two: p=0.32; paired t-tests). **C**) Analysis of behavioral responses in the two-session paradigm show that female mice show a reduced immobility responses to the tone across the first extinction session (2-way repeated measures ANOVA with multiple comparisons; p=0.01, 0.005, 0.003, and 0.01 for blocks 1-4). However, male and female mice show equivalent rates of immobility responses to the tone during the second extinction session (two-way ANOVA, p>0.8 for each block), during extinction memory retrieval (unpaired t-test, p=0.46), and at renewal (unpaired t-test, p=0.59). Male and female mice both discriminated between retrieval and renewal contexts (retrieval vs. renewal: paired t-tests, n=18 males, p=0.002; n=14 females, p=0.006).

**Extended Figure 2.**
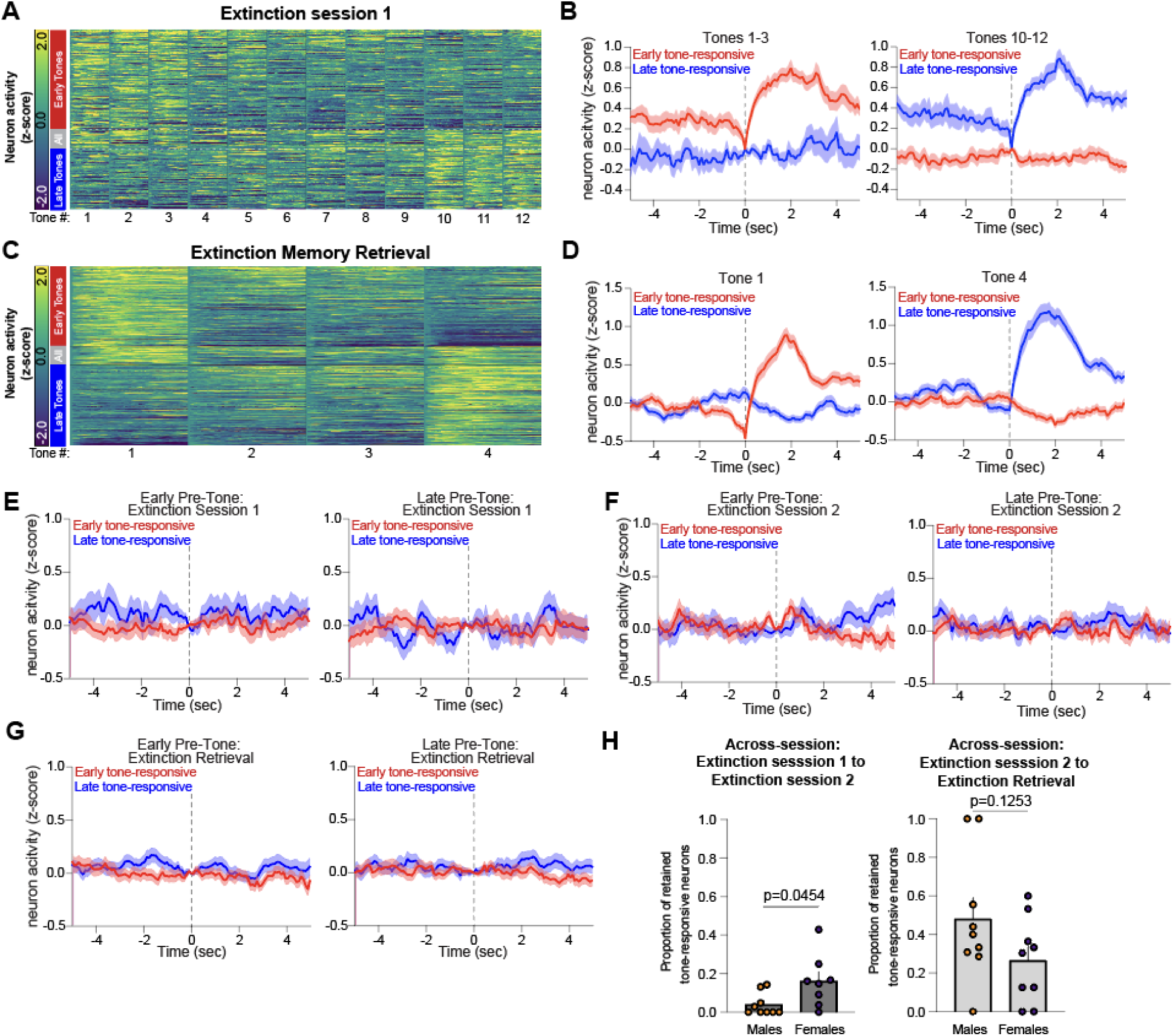
Tone-responsive neuron population dynamics within and across extinction sessions and at extinction memory retrieval. **A**) Heatmap of z-scored Ca^2+^ transients from tone-responsive neurons during the first extinction session (n=319 total neurons from 18 animals: early tone-responsive=179 neurons, all tone-responsive=24 neurons, late tone-responsive =116 neurons). **B**) Mean z-scored traces from blocks of tones from (A). **C**) Heatmap of z-scored traces from tone-responsive cells during extinction memory retrieval (n=595 total neurons from 18 animals: early tone-responsive=266 neurons, all tone-responsive=62 neurons, late tone-responsive=267 neurons). **D**) Mean z-scored traces from blocks of tones from (C). **E-G**) Mean traces of early and late activity during the first 120 seconds of each behavior session, demonstrating minimal activity of neurons subsequently assigned to early tone-responsive and late tone-responsive classifications. **H**) Female mice retain a greater proportion of tone-responsive neurons between extinction sessions 1 and 2 than male mice (left), but this effect i absent between extinction session 2 to extinction retrieval (Welch’s t-test, n=9 males and 9 females, p=0.0452 from extinction session 1 to extinction session 2; p=0.1253 from extinction session 2 to retrieval).

**Extended Figure 3.**
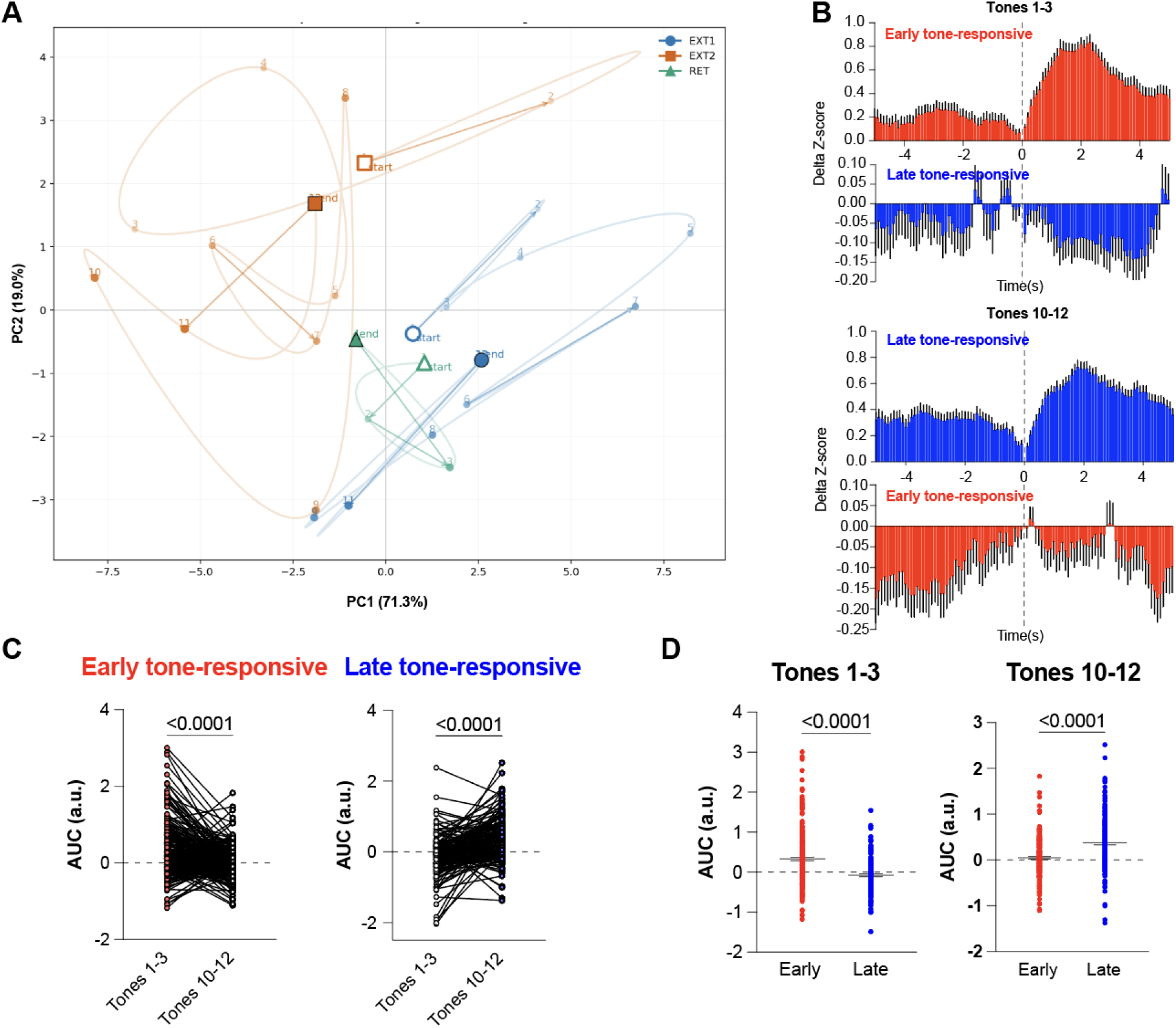
Population-state dynamics during the second extinction session. **A**) PCA trajectory plot of neuronal activity patterns during each extinction session and at extinction memory retrieval reveals the second extinction session has the longest path length and largest PC1 span (Path lengths: ext1=47.98, ext2=55.61, ret=7.33; PC1 span: ext1=10.15, ext2=12.30, ret=2.53; n: ext1=319, ext2=421, ret=595). **B**) Peristimulus time histograms from early tone-responsive and late tone-responsive neurons taken from the second extinction session shown in 100 ms bins surrounding tone-onset (paired Wilcoxon with BH-FDR across bins: 87 bins significantly different for early cells (early vs late tones), 101 bins significantly different for late cells (early vs late tones); unpaired t-test early vs late cells across bins: 95 bins significantly different). **C**) Baseline corrected AUC paired t-test of early vs late tones for early tone-onset responding cells (left, n=255 cells) and late tone-onset responding cells (right, n=145 cells) show differences in activity between early and late tones with early tone-onset neurons almost exclusively responding to early tones and late tone-onset neurons almost exclusively responding to late tones. **D**) Plot comparing AUC of early and late tone-onset responding neurons showing that the activity patterns are largely non-overlapping at early and late tone-onsets (Welch’s t-test, p<0.0001).

**Extended Figure 4.**
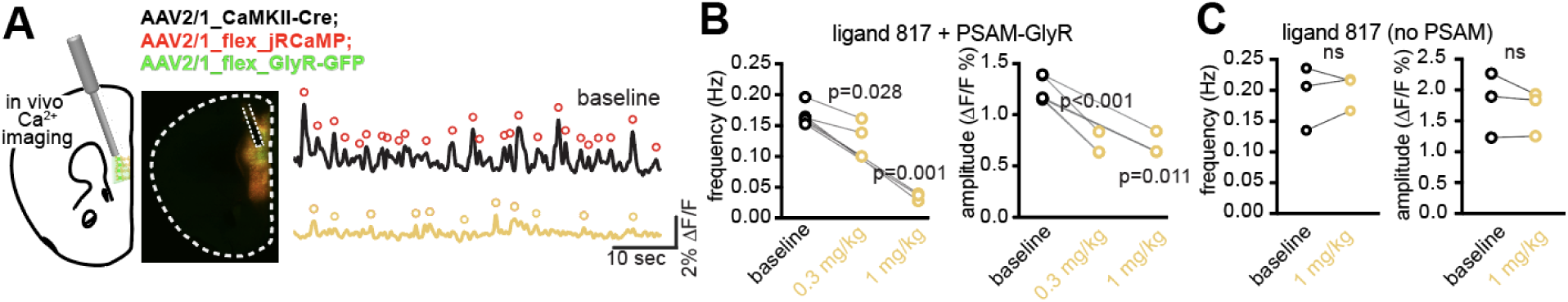
Fiber photometry recordings of PSAM-GlyR silencing of IL neuron activity from awake mice. **A**) Fiber photometry recordings were made from awake mice expressing jRCaMP and PSAM-GlyR in IL excitatory neurons (left); transients from baseline (black) and post-817 (orange) recordings were identified (see **Methods**; denoted as circles, left). **B**) 817 administration (i.p.) significantly reduced Ca^2+^ transient frequency (left: paired one-tailed t-tests, 1 mg/kg, p=0.001; 0.3 mg/kg, p=0.028) and amplitude (right: paired one-tailed t-tests, 1 mg/kg, p=0.01; 0.3 mg/kg, p=0.0002). C) Administration of 817 to mice that were not expressing the PSAM-GlyR construct did not alter transient frequency (left: paired one-tailed t-test, p=0.33) or amplitude (right; paired one-tailed t-test, p=0.18).

**Extended Figure 5.**
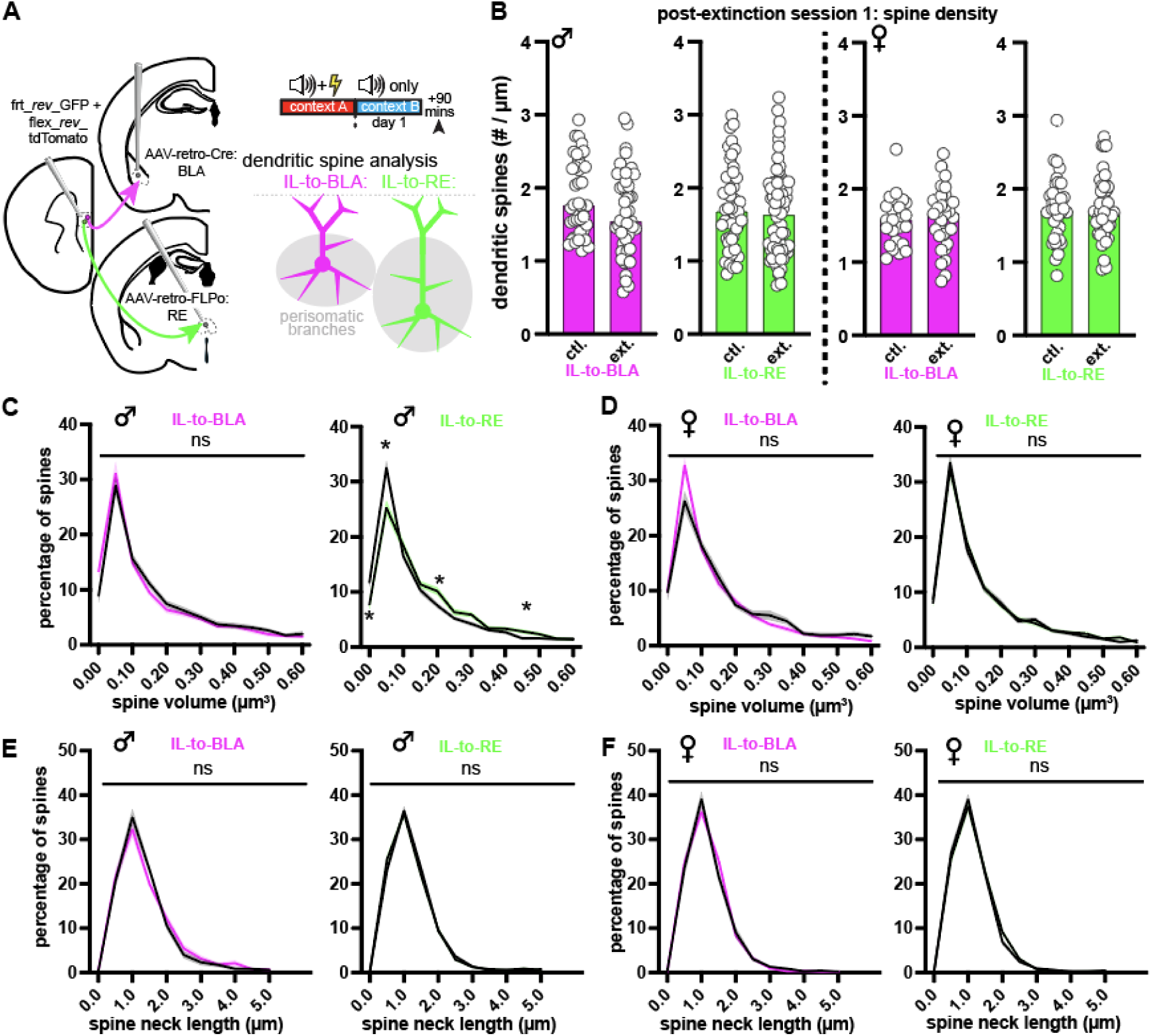
Dendritic spine morphology after a single extinction learning session. **A**) IL-to-BLA and IL-to-RE projection neurons were labeled using injections of AAVretro-Cre or AAVretro-FlpO recombinases into BLA or RE and Cre– or FlpO-dependent fluorophores into IL. Mice were sacrificed 90 mins. after the first extinction session. **B**) Dendritic spine densities were stable across perisomatic branches from both projection classes in male (left; p>0.05 for both comparisons) and female mice (right; p>0.05 for both comparisons). **C**) Spine volume distributions showed a marked stability on IL-to-BLA neurons (left; two-way ANOVA, p>0.05 for all bins) but were shifted towards larger sizes on IL-to-RE neurons in male mice (right; bin center 0, p=0.01; bin center 0.05, p=0.001, bin center 0.2, p=0.03, and bin center 0.45, p=0.04). **D**) Spine volume distributions showed no change on either projection class from female mice (two-way ANOVA, p>0.05 for all bins). **E-F**) Spine neck length distributions showed no change on either projection class from male or female mice (two-way ANOVA, p>0.05 for all bins).

**Extended Figure 6.**
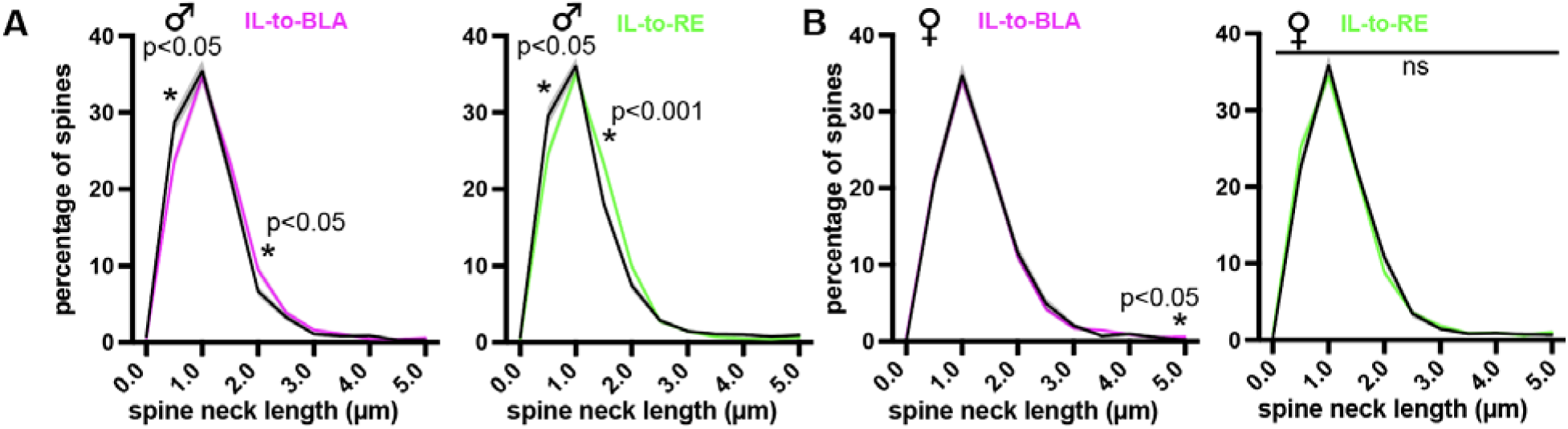
Minor modification of spine neck lengths after extinction session 2. **A**) Spine neck lengths (calculated from the center of the spine head to the center of the nearest dendritic node) from the IL-to-BLA and IL-to-RE perisomatic branches. Spine neck length distributions from mice sacrificed after the second extinction session showed minor shifts to longer necks on both projection classes in male mice (IL-to-BLA: two-way ANOVA, p=0.01 and p=0.03 at 0.5 and 2.0 bins; IL-to-RE: two-way ANOVA, p=0.01 and p<0.0001 at 0.5 and 1.5 bins). **B**) Minor or no change on either projection class from female mice IL-to-BLA: two-way ANOVA, p=0.02 at 5 µm bin; IL-to-RE: two-way ANOVA, p>0.05 for all bins).

**Extended Figure 7.**
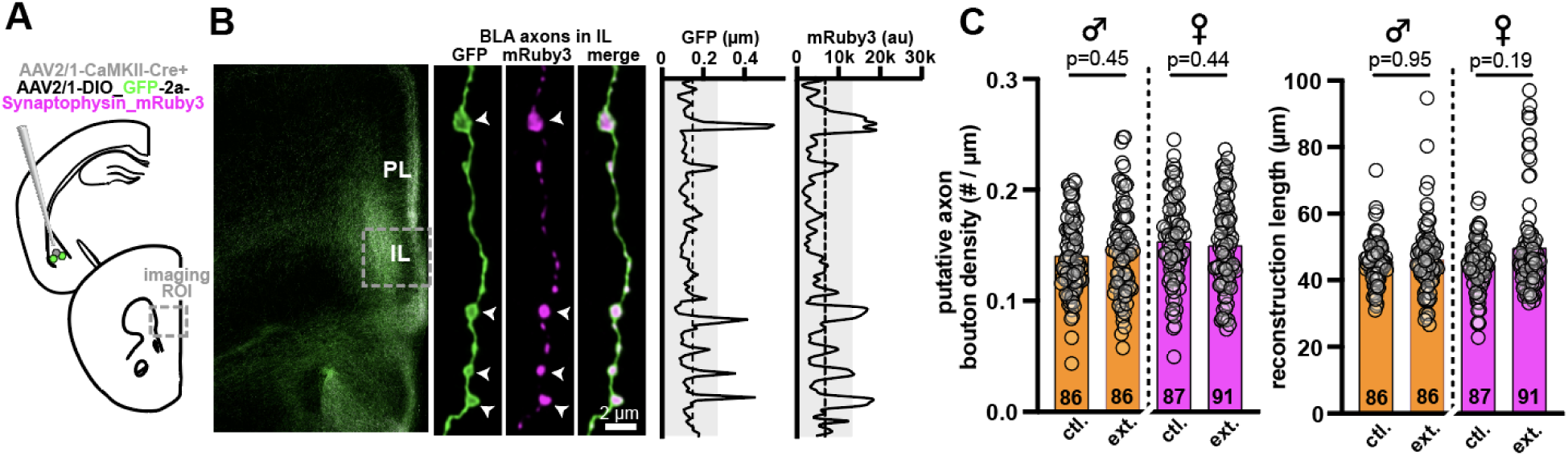
BLA axon innervation of IL is similar between male and female mice. **A**) BLA excitatory projection neurons were labeled using a mixed injection of rAAV-CaMKIIa-Cre and Cre-dependent GFP-2a-synaptiophysin_mRuby3. Mice were sacrificed 90 mins. after the second extinction session (n=4 mice per sex/group). **B**) BLA axonal innervation of IL occur primarily in the deeper layers relative to prelimbic (PL, dorsal to IL). Confocal z-stacks containing the GFP and mRuby3 signal were acquired of deeper-layer axon segments. Individual axon segments were reconstructed to analyze the cable node diameters, and putative boutons were identified as excursions > 1.5 standard deviation from the mean cable diameter. Identified boutons (arrowheads) also contained synaptophysin_mRuby signal. **C**) Bouton density was not modified by extinction learning or by sex (left; individual axon numbers denoted in the bars); reconstructed segments were ∼50 µm in length (right).

**Extended Figure 8.**
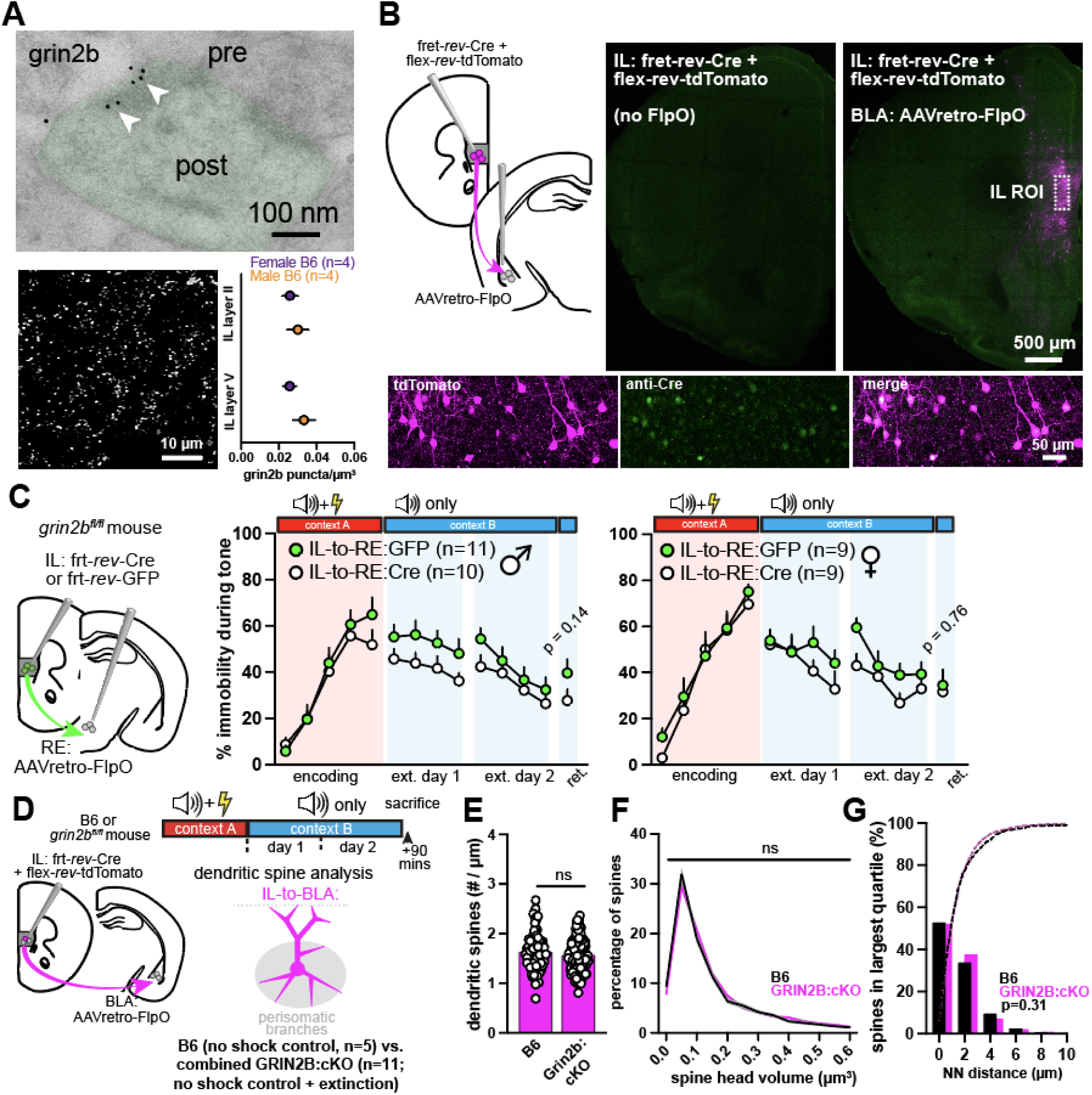
Synaptic GRIN2B localization, deletion in IL-to-RE neurons, and effects on spines relative to control mice. **A**) Postembed immunogold localization of GRIN2B to the postsynaptic density of dendritic spine synapses in IL (top). The density of immunofluorescent GRIN2B puncta in IL did not differ between male and female mice across superficial or deep cortical layers (bottom; unpaired t-test, p>0.05 for both comparisons). **B**) Dual recombinase strategy to drive FlpO-dependent Cre recombinase strictly within IL-to-BLA projection neurons (top left). Mice co-injected with frt-*rev*-Cre + flex-*rev*-tdTomato showed no tdTomato reporter expression in IL (top center), while injection of the same constructs along with AAVretro-FlpO in BLA drove IL tdTomato expression, confirming minimal leak in our Cre-dependent reporter labeling strategy (top right; tdTomato and Cre immunolabeling shown in bottom panels). **C**) Deletion of *grin2b* from IL-to-RE neurons showed no difference in extinction learning or extinction memory retrieval in male (left; two-way ANOVA for encoding, extinction session 1, and extinction session 2 where p>0.3 for all bins; unpaired t-test for retrieval, p>0.1) or female mice (right; two-way ANOVA for encoding, extinction session 1, and extinction session 2 where p>0.09 for all bins; unpaired t-test for retrieval, p>0.7). **D**) Comparison of spines after deletion of *grin2b* in male mice to spines in control B6 male mice receiving the same injections. In these analyses, *grin2b^fl/fl^*mice that received paired shocks or no-shocks during encoding were combined (see Fig. 4) and compared to no-shock B6 control mice. **E**) Spine densities on the perisomatic branches of IL-to-BLA neurons were unchanged (unpaired t-test, p>0.05). **F**) Spines from *grin2b^fl/fl^*mice showed no change in spine sizes (two-way ANOVA, p>0.05 within each bin), though this effect was small enough that there was no increase in the proportion of spines in any of the larger size bins. **G**) Deletion of *grin2b* did not change clustering of large spines along dendritic segments (Fisher’s exact test, p=0.31; Ripley’s 1D clustering analysis, p=0.9182, data not shown).

