## Supplemental Statistical Tables for "Sex-specific frontal cortical circuit mechanisms mediating fear extinction"

**SUPPLEMENTATY STATISTICAL TABLES**

| **Figure Panel** | Comparison | Test results | Significance |
|---|---|---|---|
| **Fig. 1C** | Early tone responsivity, males vs. females, extinction session 2, class proportions by sex | Male v. Female: Fischer's exact test | p<0.0001 |
| **Fig. 1I** | Per-animal tone-responsive retained proportion, comparing Ext1>Ext2 vs. Ext2>Ret within matched animals | Paired t-test<br>t(17)=3.851 | p=0.0011 |
| **Fig. 2C** | Male IL-to-BLA GFP vs. GlyR | Encoding:<br>mixed model two-way ANOVA<br>phase: F(4, 145)=60.62<br>condition: F(1, 145)=6.687<br>interaction: F (4, 145) = 1.695<br>all posthoc tests p>0.05<br><br>Extinction Day 1:<br>mixed model two-way ANOVA<br>phase: F(3, 116) = 0.56<br>condition: F(1, 116)=0.85<br>interaction: F(3, 116)=0.56<br>all posthoc tests p>0.05<br><br>Extinction Day 2:<br>mixed model two-way ANOVA<br>phase: F (3, 116) = 6.17<br>condition: F (1, 116) = 2.056<br>interaction: F (3, 116) = 0.31<br>all posthoc tests p>0.05<br><br>Extinction Retrieval: two-tailed<br>unpaired t-test<br>t(29)=1.482 | <br><br>p<0.001<br>p=0.01<br>p=0.15<br><br><br><br><br>p=0.63<br>p=0.35<br>p=0.63<br><br><br><br><br>p=0.0006<br>p=0.15<br>p=0.81<br><br><br><br>p=0.14 |
| **Fig. 2C** | Male IL-to-BLA GFP vs. GlyR<br>GFP: Ext. 2/Retrieval<br>GlyR Ext. 2/Retrieval | Paired t-tests<br>t(14)=1.038<br>t(15)=3.004 | <br>p=0.31<br>p=0.008 |
| **Fig. 2D** | Female IL-to-BLA GFP vs. GlyR | Encoding:<br>mixed model two-way ANOVA<br>phase: F (4, 166) = 38.74<br>condition: F (1, 166) = 0.24<br>interaction: F (4, 166) = 2.60<br>all posthoc tests p>0.05<br><br>Extinction Day 1: two-way ANOVA<br>phase: F (3, 136) =1.69<br>condition: F (1, 136) = 0.35<br>interaction: F (3, 136) = 0. 81<br>all posthoc tests p>0.05<br><br>Extinction Day 2: two-way ANOVA<br>phase: F (3, 136) = 2.61<br>condition: F (1, 136) = 0.65<br>interaction: F (3, 136) = 0.08<br>all posthoc tests p>0.05<br><br>Extinction Retrieval: two-tailed<br>unpaired t-test | <br><br>p<0.001<br>p=0.42<br>p=0.03<br><br><br><br>p=0.17<br>p=0.55<br>p=0.48<br><br><br><br>p=0.05<br>p=0.41<br>p=0.96 |

| | | | |
|---|---|---|---|
| | | t(34)=0.21 | p=0.82 |
| **Fig. 2D** | Female IL-to-BLA GFP vs. GlyR<br>GFP: Ext. 2/Retrieval<br>GlyR: Ext. 2/Retrieval | Paired t-tests<br>t(15)=0.60<br>t(19)=0.40 | <br>p=0.55<br>p=0.68 |
| **Fig. 2E** | Male IL-to-RE GFP vs. GlyR | Encoding:<br>mixed model two-way ANOVA<br>phase: F (4, 130) = 101.4<br>condition: F (1, 130) = 5.07<br>interaction: F (4, 130) = 0.30<br>all posthoc tests p>0.05<br><br>Extinction Day 1: two-way ANOVA<br>phase: F (3, 104) =1.99<br>condition: F (1, 104) = 4.15<br>interaction: F (3, 104) = 0.51<br>all posthoc tests p>0.05<br><br>Extinction Day 2: two-way ANOVA<br>phase: F (3, 104) =6.23<br>condition: F (1, 104) = 0.47<br>interaction: F (3, 104) = 0.08<br>all posthoc tests p>0.05<br><br>Extinction Retrieval: unpaired t-test<br>t(26)=0.43 | <br><br>p<0.001<br>p=0.02<br>p=0.87<br><br><br><br>p=0.12<br>p=0.04<br>p=0.67<br><br><br><br>p=0.0006<br>p=0.49<br>p=0.97<br><br><br><br>p=0.66 |
| **Fig. 2E** | Male IL-to-RE GFP vs. GlyR<br>GFP: Ext. 2/Retrieval<br>GlyR:Ext. 2/Retrieval | Paired t-tests<br>t(12)=0.46<br>t(14)=0.28 | <br>p=0.64<br>p=0.78 |
| **Fig. 2F** | Female IL-to-RE GFP vs. GlyR | Encoding:<br>mixed model two-way ANOVA<br>phase: F (4, 99) =95.58<br>condition: F (1,99) = 0.455<br>interaction: F (4,99) = 1.897<br>all posthoc tests p>0.05<br><br>Extinction Day 1: two-way ANOVA<br>phase: F (3, 80) =3.47<br>condition: F (1, 80) = 0.25<br>interaction: F (3, 80) = 1.35<br>all posthoc tests p>0.05<br><br>Extinction Day 2: two-way ANOVA<br>phase: F (3, 80)=5.64<br>condition: F (1, 80) = 0.16<br>interaction: F (3, 80) = 0.70<br>all posthoc tests p>0.05<br><br>Extinction Retrieval: unpaired t-test<br>t(20)=0.92 | <br><br>p<0.001<br>p=0.50<br>p=0.11<br><br><br><br>p=0.01<br>p=0.61<br>p=0.26<br><br><br><br>p=0.001<br>p=0.68<br>p=0.55<br><br><br><br>p=0.36 |
| **Fig. 2F** | Female IL-to-RE GFP vs. GlyR<br>GFP:Paired Ext. 2/Retrieval<br>GlyR:Paired Ext. 2/Retrieval | Paired t-tests<br>t(9)=0.94<br>t(11)=0.54 | <br>p=0.37<br>p=0.59 |
| **Fig. 3B** | Male IL-to-BLA Spine Density Day 2<br><br>Male IL-to-RE Spine Density Day 2 | Male IL-to-BLA: unpaired t-test<br>U=635, $n_1$=45, $n_2$=50<br>Male IL-to-RE: unpaired t-test<br>U=1342, $n_1$=60, $n_2$=59 | <br>p=0.0002<br><br>p=0.02 |

| | | | |
|---|---|---|---|
| | Female IL-to-BLA Spine Density Day 2<br><br>Female IL-to-RE Spine Density: Day 2 | Female IL-to-BLA: unpaired t-test<br>U=529, $n_1$=38, $n_2$=53<br>Female IL-to-RE: unpaired t-test<br>U=1370, $n_1$=41, $n_2$=68 | <br>p<0.0001<br><br>p=0.88 |
| **Fig. 3C** | Male IL-to-BLA Spine Volumes: Day 2<br><br><br><br><br><br><br><br>Male IL-to-RE Spine Volumes: Day 2 | Male IL-to-BLA spines:<br>two-way ANOVA (bin matching)<br>size bin: F (4.379, 422.2) =363.3<br>condition: F (1, 1157) = $3.3e^{-029}$<br>interaction: F (12, 1157) = 5.96<br>posthoc tests:<br>0.05 size bin, control vs. extinction<br>0.45 size bin, control vs. extinction<br>Male IL-to-RE spines:<br>two-way ANOVA (bin matching)<br>size bin: F (3.349, 420.9) =517.2<br>condition: F (1, 1508) = $2.7e^{-029}$<br>interaction: F (12, 1508) = 5.81<br>all posthoc tests p>0.05 | <br><br>p<0.0001<br>p>0.99<br>p<0.0001<br><br>p=0.007<br>p=0.01<br><br><br>p<0.0001<br>p>0.99<br>p<0.0001 |
| **Fig. 3D** | Male IL-to-BLA Nearest Neighbor<br>Male IL-to-RE Nearest Neighbor | Control vs. extinction: Fisher's exact test<br>Control vs. extinction: Fisher's exact test | p=0.02<br>p=0.008 |
| **Fig. 3E** | Male IL-to-BLA Ripley<br>Male IL-to-RE Ripley | Control vs. extinction: K-S test: D=0.28<br>Control vs. extinction: K-S test: D=0.25 | p=0.027<br>p=0.051 |
| **Fig. 3F** | Female IL-to-BLA Spine Volumes<br>Female IL-to-RE Spine Volumes | Female IL-to-BLA spines:<br>two-way ANOVA (bin matching)<br>size bin: F (3.787, 365.1) =348.4<br>condition: F (1, 1157) = $1.3e^{-029}$<br>interaction: F (12, 1157) = 2.65<br>all posthoc tests p>0.05<br><br>Female IL-to-RE spines:<br>two-way ANOVA (bin matching)<br>size bin: F (4.015, 456.8) =612<br>condition: F (1, 1365) = 0<br>interaction: F (12, 1365) = 1.629<br>all posthoc tests p>0.05 | <br><br>p<0.0001<br>p>0.99<br>p=0.001<br><br><br><br><br>p<0.0001<br>p>0.99<br>p=0.07 |
| **Fig. 3G** | Female IL-to-BLA Nearest Neighbor<br>Female IL-to-RE Nearest Neighbor | Control vs. extinction: Fisher's exact test<br>Control vs. extinction: Fisher's exact test | p=0.19<br>p=0.50 |
| **Fig. 3H** | Male IL-to-BLA Ripley<br>Male IL-to-RE Ripley | Control vs. extinction: K-S test: D=0.10<br>Control vs. extinction: K-S test: D=0.16 | p=0.96<br>p=0.44 |
| **Fig. 4B** | IL-to-BLA *grin2b* deletion<br>IL-to-RE *grin2b* deletion | IL-to-BLA: unpaired t-test<br>U=77, $n_1$=34, $n_2$=30<br>IL-to-RE: unpaired t-test<br>U=446.5, $n_1$=32, $n_2$=81 | <br>p<0.0001<br><br>p<0.0001 |
| **Fig. 4C left** | Male IL-to-BLA *grin2b* deletion | Encoding:<br>mixed model two-way ANOVA<br>phase: F (4, 190) =36.12<br>condition: F (1,190) = 2.53<br>interaction: F (4,190) = 0.51<br>all posthoc tests p>0.05<br><br>Extinction Day 1: two-way ANOVA<br>phase: F (3, 152) =5.32<br>condition: F (1, 152) = 6.29<br>interaction: F (3, 152) = 0.20 | <br><br>p<0.001<br>p=0.11<br>p=0.72<br><br><br><br>p=0.001<br>p=0.01<br>p=0.89 |

| | | | |
|---|---|---|---|
| | | all posthoc tests p>0.05<br><br>Extinction Day 2: two-way ANOVA<br>phase: F (3, 152) =8.99<br>condition: F (1, 152) = 25.65<br>interaction: F (3, 152) = 0.46<br>posthoc tests:<br>block 1, GFP vs. Cre<br>block 2, GFP vs. Cre<br>block 3, GFP vs. Cre<br>block 4, GFP vs. Cre<br><br><br>Extinction Retrieval: unpaired t-test<br>t(38)=2.19 | <br><br><br>p<0.0001<br>p<0.0001<br>p=0.70<br><br>p=0.003<br>p=0.04<br>p=0.08<br>p=0.04<br><br><br><br>p=0.03 |
| **Fig. 4C right** | Female IL-to-BLA *grin2b* deletion | Encoding:<br>mixed model two-way ANOVA<br>phase: F (4, 200) =35.83<br>condition: F (1,200) = 0.79<br>interaction: F (4,200) = 0.23<br>all posthoc tests p>0.05<br><br>Extinction Day 1: two-way ANOVA<br>phase: F (3, 160) =5.63<br>condition: F (1, 160) = 1.33<br>interaction: F (3, 160) = 0.54<br>all posthoc tests p>0.05<br><br>Extinction Day 2: two-way ANOVA<br>phase: F (3, 160)=7.07<br>condition: F (1, 160) = 7.47<br>interaction: F (3, 160) = 0.93<br>all posthoc tests p>0.05<br><br>Extinction Retrieval: unpaired t-test<br>t(40)=1.56 | <br><br>p<0.001<br>p=0.37<br>p=0.92<br><br><br><br>p=0.001<br>p=0.25<br>p=0.65<br><br><br><br>p=0.0002<br>p=0.007<br>p=0.42<br><br><br><br>p=0.12 |
| **Fig. 4E** | Male IL-to-BLA *grin2b* deletion:<br>dendritic spine densities | Control vs. extinction: unpaired t-test<br>U=2322, $n_1$=67, $n_2$=74 | <br>p=0.51 |
| **Fig. 4F** | Male IL-to-BLA *grin2b* deletion:<br>dendritic spine volume distributions | Spine volume: two-way ANOVA<br>bin: F (12, 1807) =507.6<br>condition: F (1, 1807) = 0<br>interaction: F (12, 1807) = 1.125<br>all posthoc tests p>0.05 | <br>p<0.001<br>p=0.99<br>p=0.33 |
| **Fig. 4G** | Male IL-to-BLA *grin2b* deletion:<br>dendritic spine clustering | Control vs. extinction: Fisher's exact test | p=0.31 |
| **Extended Data 1B** | One extinction session vs.<br>two extinction sessions | Encoding:<br>two-way ANOVA<br>phase: F (4, 134) =25.54<br>condition: F (1, 134) = 0.94<br>interaction: F (4, 134) = 0.63<br>all posthoc tests p>0.05<br><br>Extinction Day 1: | <br><br>p<0.0001<br>p=0.33<br>p=0.63 |

| | | | |
|---|---|---|---|
| | | two-way ANOVA<br>phase: F (3, 108) =3.26<br>condition: F (1, 108) = 0.25<br>interaction: F (3, 108) = 0.44<br>all posthoc tests p>0.05<br><br>Retrieval: unpaired, one-tailed t-test<br>t(27)=1.75 | <br>p=0.02<br>p=0.61<br>p=0.72<br><br><br><br>p=0.04 |
| **Extended Data 1B** | Context-dependent discrimination, two extinction sessions | within mouse, paired retrieval vs. renewal: week 1<br>t(15)=5.58 | <br><br>p<0.0001 |
| **Extended Data 1B** | Context-dependent discrimination, one extinction session | within mouse, paired retrieval vs. renewal: week 1<br>t(12)=1.71 | <br><br>p=0.11 |
| **Extended Data 1B** | Context-dependent discrimination, two extinction sessions | within mouse, paired retrieval vs. renewal: week 2<br>t(15)=3.07 | <br><br>p=0.007 |
| **Extended Data 1B** | Context-dependent discrimination, one extinction session | within mouse, paired retrieval vs. renewal: week 2<br>t(12)=1.03 | <br><br>p=0.32 |
| **Extended Data 1C** | Male vs. female mice | Encoding:<br>two-way ANOVA<br>phase: F (4, 463) =184.2<br>sex: F (1, 463) = 3.09<br>interaction: F (4, 463) = 1.24<br>all posthoc tests p>0.05<br><br>Extinction Day 1: two-way ANOVA<br>phase: F (3, 376) =6.58<br>sex: F (1, 376) = 34.07<br>interaction: F (3, 376) = 0.42<br>posthoc tests:<br>block 1<br>block 2<br>block 3<br>block 4<br><br>Extinction Day 2: two-way ANOVA<br>phase: F (3, 328)=5.15<br>sex: F (1, 328) = 0.25<br>interaction: F (3, 323) = 0.33<br>all posthoc tests p>0.05<br><br>Extinction Retrieval: unpaired t-test<br>t(55)=0.74<br><br>Renewal: unpaired t-test<br>t(30)=0.53 | <br><br>p<0.0001<br>p=0.07<br>p=0.28<br><br><br><br>p=0.0002<br>p<0.0001<br>p=0.73<br><br>p=0.02<br>p=0.005<br>p=0.001<br>p=0.01<br><br><br>p=0.001<br>p=0.61<br>p=0.79<br><br><br><br>p=0.46<br><br><br>p=0.59 |
| **Extended Data 2H** | Male vs female mice, across session retention extinction 1 to extinction 2<br><br>Male vs female mice, across session retention extinction 2 to retrieval | Male vs female: unpaired, one-tailed t-test<br>t(16)=1.98<br>Male vs. female: unpaired, t-test<br>t(14)=1.63 | p=0.0454<br><br><br>p=0.1253 |
| **Extended Data 3B** | Tones 1-3 vs. Tones 10-12: Peristimulus Time Histogram for Early Tone Responding Neurons (100 ms bins) | Tones 1-3 vs Tones 10-12: paired, Wilcoxon with BH-FDR | 87 bins<br>p<0.05 |

| | | | |
|---|---|---|---|
| | Tones 1-3 vs Tones 10-12: Peristimulus Time Histogram for Late Tone Responding Neurons (100 ms bins)<br><br>Early vs Late Tone Responding Neurons: PSTH for Tones 1-3<br><br>Early vs Late Tone Responding Neurons: PSTH for Tones 10-12 | Tones 1-3 vs Tones 10-12: paired, Wilcoxon with BH-FDR<br><br><br>Early vs Late Neurons: unpaired, t-test<br><br><br>Early vs Late Neurons: unpaired, t-test | 101 bins<br>$p<0.05$<br><br><br>95 bins<br>$p<0.05$<br><br>95 bins<br>$p<0.05$ |
| **Extended Data 3C** | Tones 1-3 vs Tones 10-12: Early Tone-Responsive Neurons AUC from Extinction 2<br><br>Tones 1-3 vs Tones 10-12: Late Tone-Responsive Neurons AUC from Extinction 2 | Tones 1-3 vs Tones 10-12: paired, t-test $t(254)=5.70$<br><br>Tones 1-3 vs Tones 10-12: paired, t-test $t(162)=7.56$ | $p<0.0001$<br><br><br>$p<0.0001$ |
| **Extended Data 3D** | Early Tone-Responsive vs Late Tone-Responsive Neurons: Tones 1-3 AUC<br><br>Early Tone-Responsive vs Late Tone-Responsive Neurons: Tones 10-12 AUC | Early vs Late Neurons: unpaired, t-test $t(393)=7.16$<br><br>Early vs Late Neurons: unpaired, t-test $t(258)=5.96$ | $p<0.0001$<br><br><br>$p<0.0001$ |
| **Extended Data 4B** | Inhibition of $Ca^{2+}$ transient frequency: baseline vs. 0.3 mg/kg<br>Inhibition of $Ca^{2+}$ transient frequency: baseline vs. 1 mg/kg<br>Inhibition of $Ca^{2+}$ transient amplitude: baseline vs. 0.3 mg/kg<br>Inhibition of $Ca^{2+}$ transient amplitude: baseline vs. 1 mg/kg | One-tailed Paired t-test, $t(2)=4.052$<br><br>One-tailed Paired t-test, $t(2)=20.50$<br><br>One-tailed Paired t-test, $t(2)=49.06$<br><br>One-tailed Paired t-test, $t(2)=6.48$ | $p=0.028$<br><br>$p=0.001$<br><br>$p=0.002$<br><br>$p=0.01$ |
| **Extended Data 4C** | Inhibition of $Ca^{2+}$ transient frequency: baseline vs. 1 mg/kg (no PSAM)<br>Inhibition of $Ca^{2+}$ transient amplitude: baseline vs. 1 mg/kg (no PSAM) | One-tailed Paired t-test, $t(2)=0.509$<br><br>One-tailed Paired t-test, $t(2)=1.152$ | $p=0.33$<br><br>$p=0.18$ |
| **Extended Data 5B** | Male IL-to-BLA Spine Density Day 1<br><br>Male IL-to-RE Spine Density Day 1<br><br>Female IL-to-BLA Spine Density Day 1<br><br>Female IL-to-RE Spine Density Day 1 | Male IL-to-BLA: unpaired t-test<br>$U=774$, $n_1=41$, $n_2=49$<br>Male IL-to-RE: unpaired t-test<br>$U=1514$, $n_1=49$, $n_2=68$<br>Female IL-to-BLA: unpaired t-test<br>$U=346$, $n_1=23$, $n_2=33$<br>Female IL-to-RE: unpaired t-test<br>$U=816$, $n_1=36$, $n_2=46$ | <br>$p=0.06$<br><br>$p=0.40$<br><br>$p=0.58$<br><br>$p=0.91$ |
| **Extended Data 5C** | Male IL-to-BLA Spine Volumes: Day 1<br><br><br><br><br><br><br><br>Male IL-to-RE Spine Volumes Day 1 | Male IL-to-BLA spines:<br>two-way ANOVA (bin matching)<br>size bin: $F(3.186, 296.8.2) = 200.4$<br>condition: $F(1, 1118) = 1.2e^{-029}$<br>interaction: $F(12, 1118) = 2.012$<br>all posthoc tests $p>0.05$<br><br>Male IL-to-RE spines:<br>two-way ANOVA (bin matching)<br>size bin: $F(3.854, 455.1) =345.5$ | <br><br>$p<0.0001$<br>$p>0.99$<br>$p=0.02$<br><br><br><br><br>$p<0.0001$ |

| | | | |
|---|---|---|---|
| | | condition: F (1, 1417) = 0<br>interaction: F (12, 1417) = 10.01<br>posthoc tests:<br>0.0 size bin, control vs. extinction<br>0.05 size bin, control vs. extinction<br>0.20 size bin, control vs. extinction<br>0.45 size bin, control vs. extinction | p>0.99<br>p<0.0001<br><br>p=0.01<br>p=0.001<br>p=0.03<br>p=0.04 |
| **Extended Data 5D** | Female IL-to-BLA Spine Volumes: Day 1<br><br><br><br><br><br><br><br>Female IL-to-RE Spine Volumes: Day 1 | Female IL-to-BLA spines:<br>two-way ANOVA (bin matching)<br>size bin: F (4.317, 257.2) = 238.8<br>condition: F (1, 715) = 3.4e$^{-029}$<br>interaction: F (12, 715) = 3.754<br>all posthoc tests p>0.05<br><br><br>Female IL-to-RE spines:<br>two-way ANOVA (bin matching)<br>size bin: F (3.163, 274.1) =393.9<br>condition: F (1, 1040) = 0<br>interaction: F (12, 1040) = 0.57<br>all posthoc tests p>0.05 | <br><br>p<0.0001<br>p>0.99<br>p<0.0001<br><br><br><br><br><br>p<0.0001<br>p>0.99<br>p=0.86 |
| **Extended Data 5E** | Male IL-to-BLA Spine Neck Length: Day 1<br><br><br><br><br><br><br><br><br>Male IL-to-RE Spine Neck Length: Day 1 | Male IL-to-BLA spines:<br>two-way ANOVA (bin matching)<br>size bin: F (1.960, 187.5) = 315.6<br>condition: F (1, 957) = 0.0007<br>interaction: F (10, 957) = 1.53<br>all posthoc tests p>0.05<br><br><br><br>Male IL-to-RE spines:<br>two-way ANOVA (bin matching)<br>size bin: F (3.084, 369.8) =1065<br>condition: F (1, 1199) = 0<br>interaction: F (10, 1199) = 1.61<br>all posthoc tests p>0.05 | <br><br>p<0.0001<br>p=0.97<br>p=0.12<br><br><br><br><br><br><br>p<0.0001<br>p>0.99<br>p=0.09 |
| **Extended Data 5F** | Female IL-to-BLA Spine Neck Length: Day 1<br><br><br><br><br><br><br>Female IL-to-RE Spine Neck Length: Day 1 | Female IL-to-BLA spines:<br>two-way ANOVA (bin matching)<br>size bin: F (2.994, 181.1) = 586.6<br>condition: F (1, 605) = 9.0e$^{-029}$<br>interaction: F (10, 605) = 1.95<br>all posthoc tests p>0.05<br><br>Female IL-to-RE spines:<br>two-way ANOVA (bin matching)<br>size bin: F (2.564, 225.6) =802.4<br>condition: F (1, 880) = 0<br>interaction: F (10, 880) = 1.056<br>all posthoc tests p>0.05 | <br><br>p<0.0001<br>p>0.99<br>p=0.03<br><br><br><br><br>p<0.0001<br>p>0.99<br>p=0.39 |
| **Extended Data 6A** | Male IL-to-BLA Spine Neck Length: Day 2 | Male IL-to-BLA spines:<br>two-way ANOVA (bin matching)<br>size bin: F (2.685, 305.7) = 703.5<br>condition: F (1, 1067) = 0<br>interaction: F (10, 1067) = 4.221<br>posthoc tests: | <br><br>p<0.0001<br>p>0.99<br>p<0.0001 |

| | | | |
|---|---|---|---|
| | <br><br><br>Male IL-to-RE Spine Neck Length:<br>Day 2 | 0.5 size bin, control vs. extinction<br>2.0 size bin, control vs. extinction<br><br>Male IL-to-RE spines:<br>two-way ANOVA (bin matching)<br>size bin: F (3.080, 393) =1090<br>condition: F (1, 1276) = 0<br>interaction: F (10, 1276) = 10.13<br>posthoc tests:<br>0.5 size bin, control vs. extinction<br>1.5 size bin, control vs. extinction | p=0.01<br>p=0.03<br><br><br><br>p<0.0001<br>p>0.99<br>p<0.0001<br><br>p=0.01<br>p<0.0001 |
| **Extended Data 6B** | Female IL-to-BLA Spine Neck Length:<br>Day 2<br><br><br><br><br><br><br><br><br>Female IL-to-RE Spine Neck Length:<br>Day 2 | Female IL-to-BLA spines:<br>two-way ANOVA (bin matching)<br>size bin: F (3.064, 300) = 666.1<br>condition: F (1, 979) = 0<br>interaction: F (10, 979) = 0.28<br>posthoc tests:<br>5.0 size bin, control vs. extinction<br><br><br>Female IL-to-RE spines:<br>two-way ANOVA (bin matching)<br>size bin: F (3.506, 402.9) =808.6<br>condition: F (1, 1149) = 0.0006<br>interaction: F (10, 1149) = 1.774<br>all posthoc tests p>0.05 | <br><br>p<0.0001<br>p>0.99<br>p=0.98<br><br>p=0.02<br><br><br><br><br>p<0.0001<br>p=0.97<br>p=0.06 |
| **Extended Data S7C** | BLA-to-IL axon bouton density (#/µm):<br>Male control vs. extinction<br><br>Female control vs. extinction<br><br>BLA-to-IL axon reconstruction length | Male IL-to-BLA: unpaired t-test<br>U=3454, $n_1$=86, $n_2$=86<br>Female IL-to-BLA: unpaired t-test<br>U=3694, $n_1$=87, $n_2$=91<br><br>Male IL-to-BLA: unpaired t-test<br>U=3678, $n_1$=86, $n_2$=86<br>Female IL-to-BLA: unpaired t-test<br>U=3514, $n_1$=87, $n_2$=91 | <br>p=0.45<br><br>p=0.44<br><br><br>p=0.95<br><br>p=0.19 |
| **Extended Data S8A** | GRIN2B puncta across btween male and female mice<br>IL Layer II (#/µm$^3$)<br><br>IL LayerV (#/µm$^3$) | <br>unpaired t-test<br>U=6, $n_1$=4, $n_2$=4<br><br>unpaired t-test<br>U=4, $n_1$=4, $n_2$=4 | <br><br>p=0.68<br><br><br>p=0.34 |
| **Extended Data 8C** | Male IL-to-RE *grin2b* deletion | Encoding:<br>mixed model two-way ANOVA<br>phase: F (4, 95) =31.75<br>condition: F (1, 95) = 1.011<br>interaction: F (4, 95) = 0.55<br>all posthoc tests p>0.05<br><br>Extinction Day 1: two-way ANOVA<br>phase: F (3, 76) = 1.029<br>condition: F (1, 76) = 8.687<br>interaction: F (3, 76) = 0.02<br>all posthoc tests p>0.05 | <br><br>p<0.001<br>p=0.31<br>p=0.69<br><br><br><br>p=0.38<br>p=0.004<br>p=0.99 |

| | | | |
|---|---|---|---|
| | Female IL-to-RE *grin2b* deletion | Extinction Day 2: two-way ANOVA<br>phase: F (3, 76) = 5.285<br>condition: F (1, 76) = 3.606<br>interaction: F (3, 76) = 0.22<br>all posthoc tests p>0.05<br><br>Extinction Retrieval: unpaired t-test<br>t(19)=1.523<br><br>Encoding:<br>mixed model two-way ANOVA<br>phase: F (4, 80) =34.90<br>condition: F (1, 80) = 0.88<br>interaction: F (4,80) = 0.28<br>all posthoc tests p>0.05<br><br>Extinction Day 1: two-way ANOVA<br>phase: F (3, 64) = 1.884<br>condition: F (1, 64) = 1.889<br>interaction: F (3, 64) = 0.54<br>all posthoc tests p>0.05<br><br>Extinction Day 2: two-way ANOVA<br>phase: F (3, 64) =4.617<br>condition: F (1, 64) = 7.052<br>interaction: F (3, 64) = 0.55<br>all posthoc tests p>0.05<br><br>Extinction Retrieval: unpaired t-test<br>t(16)=0.30 | <br>p=0.002<br>p=0.06<br>p=0.88<br><br><br><br>p=0.14<br><br><br><br>p<0.001<br>p=0.35<br>p=0.88<br><br><br><br>p=0.14<br>p=0.17<br>p=0.65<br><br><br><br>p=0.005<br>p=0.01<br>p=0.64<br><br><br><br>p=0.76 |
| **Extended Data 8E** | Male IL-to-RE B6 vs. *grin2b* deletion: Spine density | Male IL-to-BLA B6 vs. *grin2b*-deleted: unpaired t-test<br>U=4697, $n_1$=74, $n_2$=141 | p=0.23 |
| **Extended Data 8F** | Male IL-to-RE B6 vs. *grin2b* deletion: Spine head volume | Male IL-to-BLA B6 vs. *grin2b*-deleted: two-way ANOVA (bin matching)<br>size bin: F (4.627, 1063) = 707.4<br>condition: F (1, 2756) = 0<br>interaction: F (12, 2756) = 2.809<br>all posthoc tests p>0.05 | <br><br>p<0.0001<br>p>0.99<br>p=0.0008 |
| **Extended Data 8G** | Male IL-to-RE B6 vs. *grin2b* deletion: Nearest neighbor | B6 vs. *grin2b*-deleted: Fisher's exact test | p=0.31 |
